# Fibroblast mechanoperception instructs pulmonary developmental and pattern specification gene expression programs

**DOI:** 10.1101/2024.12.26.630418

**Authors:** Andrew E. Miller, Ping Hu, Rishi Bhogaraju, Riley T. Hannan, Daniel Abebayehu, Mete Civelek, Thomas H. Barker

**Affiliations:** Department of Biomedical Engineering, Schools of Engineering and Medicine, University of Virginia, Charlottesville, VA 22908; Division of Pulmonary & Critical Care Medicine, School of Medicine, University of Virginia, Charlottesville, VA 22908; Department of Genome Sciences, School of Medicine, University of Virginia, Charlottesville, VA 22908

## Abstract

Dysregulation of the cellular mechanisms that coordinate the interpretation and transduction of microenvironmental biophysical signals are a unifying feature of tissue remodeling pathologies such as fibrosis and cancer. While genomic regulation downstream of normal mechanotransduction (i.e. cases where cells sense soft and stiff appropriately) is well studied, significantly less is known about the consequences of abnormal mechanoperception and subsequent misinterpretation of the mechanical environment. Leveraging Thy-1 (a.k.a. CD90) loss as a model of impaired mechanoperception, we employed ATAC- and RNA-sequencing in parallel to characterize the changes in lung fibroblast genomic activity in response to a combination of substrate stiffness and culture time. Notably, we find perturbed mechanoperception elicits a near-complete shutdown of *HOXA5*, a transcription factor responsible for pattern specification and development in the nascent lung. *In vitro* investigation of HOXA5 expression reveals a potential mechanism connecting increased α_v_ integrin signaling, cytoskeletal tension, and SRC kinase activity to HOXA5 silencing. These results establish novel links between integrin signaling and the expression dynamics of genes necessary for tissue formation and regeneration in the injured and/or developing lung, particularly *HOXA5*.

## Introduction

Appropriate interpretation and transmission of biophysical signals from the extracellular microenvironment is essential to the maintenance of organism homeostasis both during development and in adult life. The orchestrated processes and componentry utilized by cells to transmit such physical forces is termed mechanotransduction. In adult tissues, mechanotransduction coordinates tissue repair and extracellular matrix (ECM) turnover in the wake of tissue injury and as a part of normal tissue homeostasis. Perhaps paradoxically, normal mechanotransduction can passively contribute to pathogenesis as a consequence of gradual ECM stiffening and subsequent elevations in signaling flux through mechanically responsive signaling pathways. In contrast, disease initiation from a homeostatic, soft tissue state requires a combination of soluble cues (e.g. transforming growth factor β, connective tissue growth factor, IL-1β, and TNF-α) and aberrant cellular responses to normal mechanical cues arising from dysregulation of the mechanotransduction machinery.^1–4^ The latter implies a causal role for perturbed mechanoperception – the cell’s ability to appropriately interpret, and respond to, the biophysical cues in its surrounding microenvironment – in the induction of pathologies characterized by deviations from normal ECM architecture and mechanics such as fibrosis and arthritis. In this manuscript, we define *mechanotransduction* as the innate mechanisms used by the cell to sense and convert mechanical cues into discernable signals that instruct function. In juxtaposition, *mechanoperception* refers to the machinery and processes that enable the cell to appropriately interpret such mechanical cues. It is worth noting that these terms are not mutually exclusive from one another.

At the molecular level, integrin receptor engagement of their cognate ECM ligand(s) (e.g. fibronectin (Fn), collagens, laminin) catalyzes the formation of the mechanotransduction signaling apparatus by promoting the recruitment of protein scaffolding to the intracellular domains of the integrin toward the creation of integrin adhesion complexes (IAC), or focal adhesions (FA), which physically tether the cell-adjacent ECM to the nucleus via the actin cytoskeleton. The recruitment of protein kinases during FA formation and maturation (e.g. ILK, PAK, and the SRC family kinases (SFKs)) generates a robust biochemical signaling hub that links integrin force transmission to the activation of cytosolic biochemical signaling cascades including: PI3K/AKT, WNT/β-Catenin, and MAPK.^5–7^ In fibrotic pathologies, aberrant activity of these pathways is a common disease signature and promotes the formation of persistent, contractile myofibroblasts (MFs) that serve as the main drivers of scar formation and organ dysfunction.^8^

Thy-1, a.k.a. CD90, is a glycophosphatidylinositol (GPI)-anchored protein located on the outer leaflet of the plasma membrane. Canonically described as an immune and mesenchymal cell marker, Thy-1 dysregulation is now tied to a wide range of pathologies including fibrotic disorders, cancers, and neurological diseases.^9–14^ Mechanistically, Thy-1 plays a direct role in cell mechanoperception and normal mechanotransduction by regulating integrin affinity for ECM as well as facilitating the co-clustering of FA-bound kinases and kinase inhibitors that confer normal mechanosignaling in response to force application on FAs.^15^ In pulmonary fibroblasts, Thy-1 has been shown to act as a stiffness-dependent inhibitor of integrin α_v_ activation by weakly restraining membrane-bound integrin α_v_ in a closed, inactive confirmation on compliant, physiologically “normal” ECM stiffnesses. The inhibitory activity of Thy-1 is naturally overcome in stiff environments through inside-out activation of integrin α_v_ to maintain normal mechanotransduction in response to increases in the tensile strength of the surrounding ECM. When Thy-1 expression is lost due to epigenetic silencing or cell surface shedding, fibroblasts display noticeably elevated integrin α_v_ activation and concomitantly assume an apoptotic-resistant, myofibroblast-like state in soft environments that typically inhibit myofibroblast activation.^15, 16^ Thus, loss of Thy-1 disrupts stiffness-dependent regulation of integrin α_v_ surface activation and leads to divergent interpretation of the mechanical properties of the ECM. This, collectively, implies a direct role for Thy-1 in the maintenance of canonical fibroblast mechanoperception.

In opposition to the tissue destructive phenotypes that emerge in fibrosis, tissue regeneration engages well-regulated programs of tissue/organ formation that direct the developing structures in the embryo.^17–19^ Thus, adult cells’ ability to properly engage developmental programs is likely a key element of successful tissue regeneration. In development, the homeobox (HOX) family of transcription factors collectively establish tissue patterning along the anterior-posterior (i.e. head-to-tail) axis. With respect to the developing lung, the *HOX5* genes, predominately *HOXA5*, are required for complete airway maturation.^20, 21^ Homozygous knockout (KO) of *HOXA5* induces an embryonic lethal phenotype underscored by alveolar simplification and tracheal malformation.^22^ Heterozygous *HOXA5* KO mice are viable but display significantly impaired alveologenesis, mislocalization of pulmonary myofibroblasts, and aberrant organization of the pulmonary elastin network.^23–25^ In adult tissues, *HOXA5* silencing in non-small cell lung cancer fibroblasts has been shown to dysregulate cytoskeletal organization and enhance cell spreading through elevated ARP2/3 levels.^26^ Conversely, forced overexpression of HOXA5 in hypertrophic scar-derived myofibroblasts significantly reduces their contractility and collagen secretion.^27^ While the downstream impacts of HOXA5 dysregulation suggest a regulatory role in fibroblast mechanotransduction, a role for ECM mechanics and integrin signaling in the upstream control of HOXA5 expression remains largely unknown.

Here, we leveraged the expression of Thy-1 to drive normal or aberrant mechanoperception phenotypes in human lung fibroblasts. Using combinatorial multi-omics (ATAC- and RNA-) sequencing we developed an atlas of chromatin accessibility and transcriptome expression in human lung fibroblasts across gradients of both time and substrate stiffness to determine how normal and abnormal fibroblast mechanotransduction may lead to long-term reprogramming of cell phenotypes through epigenetic regulation of chromatin structure and transcript expression. We also sought to explore whether misinterpretation of ECM mechanics by fibroblasts would yield a predictable genomic and transcriptomic output based on the cytoskeletal phenotypes we have previously reported in the Thy-1 negative phenotype.

## Results

### Thy-1 silencing perturbs lung fibroblast mechanosensing

We first sought to develop a persistent model of Thy-1 inhibition for our studies. To this end, we employed CRISPR-Cas9 to generate Thy-1^KO^ fibroblasts using a commercially available immortalized human lung fibroblast cell line (iHLF). Successful knockout of Thy-1 on the cell surface was confirmed for all clones via flow cytometry (Figure S1). *In vitro* analysis of the Thy-1^KO^ mechanical phenotype aligned well with previous observations in primary mouse and human lung fibroblasts as well as established Thy-1^+^ and Thy-1^-^ fibroblast cell lines.^15, 28^ As expected, Thy-1^KO^ fibroblasts cultured on soft (2kPa), Fn-coated polyacrylamide hydrogels exhibited enhanced cell spreading compared to their WT counterparts as well as defects in their capacity to further spread in response to increasing substrate stiffness (Figure 1A, B). Given that Thy-1 acts as a weak inhibitor of integrin α_v_β_3_ avidity, we next assessed the stoichiometry of the canonical fibroblast Fn-binding integrins, α_v_β_3_ and α_5_β_1_, following Thy-1 silencing. Indeed, loss of Thy-1 was sufficient to promote a significant increase in the ratio of active β_3_:β_1_ integrin on the fibroblast cell surface independent of stiffness (Figure 1C). Interestingly, the impact of Thy-1 loss on β_3_:β_1_ activation levels was most pronounced on soft hydrogels and displayed a stiffness-dependent decrease which agrees with our past findings and the established literature illustrating the dominant role of integrin β_1_ (α_5_β_1_) in supporting the high tensile forces generated across FAs in fibroblasts cultured on rigid substrates.^15, 29^ Paxillin staining and quantification of FA area did not illustrate a similar, stiffness-dependence, but a significant elevation in FA size was observed when culturing Thy-1^KO^ fibroblasts on soft substrates (Figure 1D). The acquisition of a well-spread, activated phenotype on soft ECM and an inability to continue spreading in response to elevated ECM rigidity are conserved features of the Thy-1 negative fibroblast. This is accompanied by a notable shift in integrin usage away from α_5_β_1_ towards α_v_β_3_ which, based on the established literature, will have a significant impact on fibroblast behavior by disrupting the physiologically normal patterns of integrin usage at a given stiffness.^28, 30^

**Figure 1.**
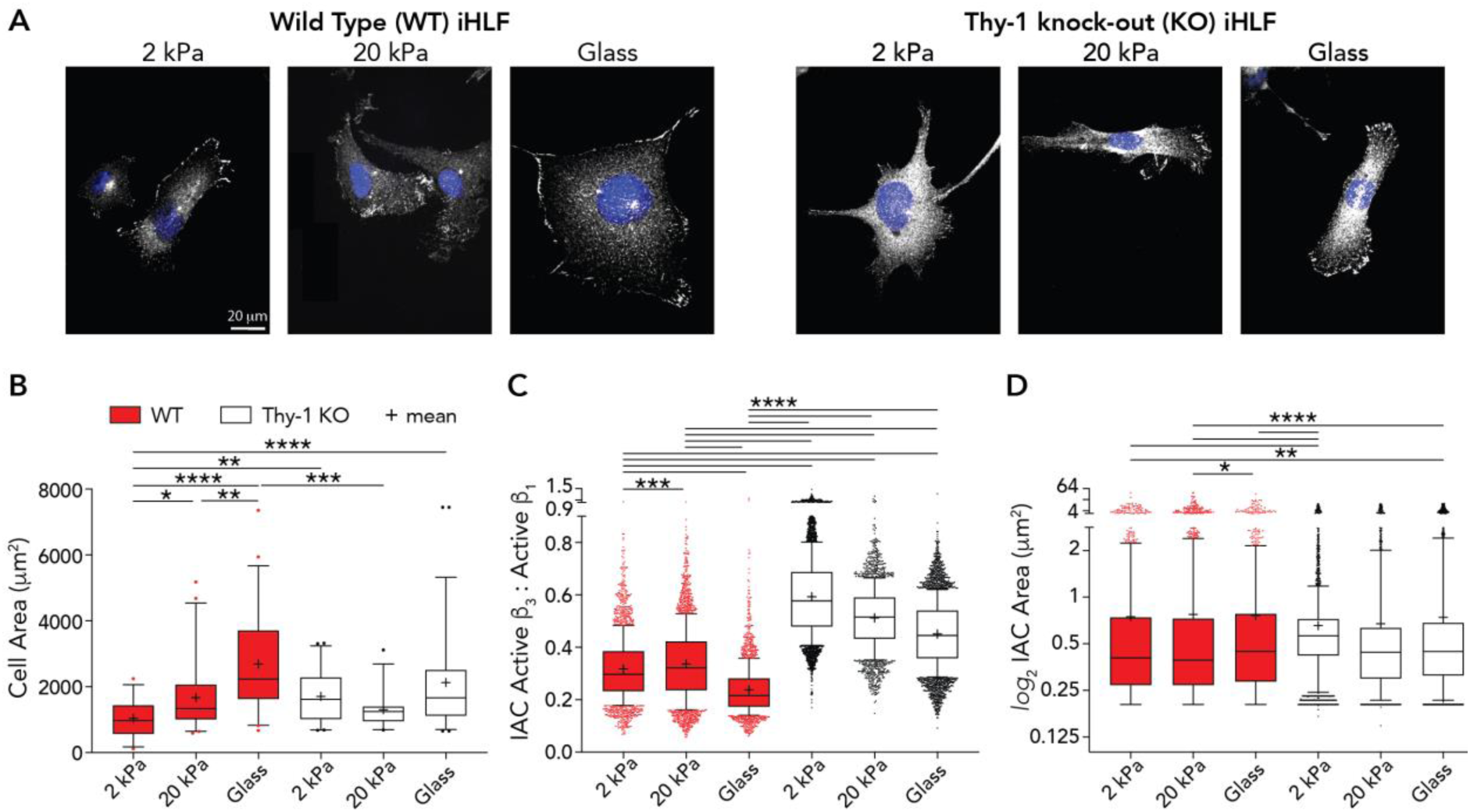
Loss of Thy-1 impairs fibroblast mechanosensing, integrin stoichiometry, and focal adhesion dynamics. (A) Representative images of aberrant cell spreading behavior following *Thy-1* knockout. Thy-1^KO^ fibroblasts present with large focal adhesion (FA) complexes and a well spread phenotype on fibronectin-coated soft hydrogels that remains unchanged with increasing substrate stiffness (Scale bar = 20μm; Blue = DAPI; White = Paxillin). (B) Cell area quantification as a function of substrate stiffness between WT and Thy-1^KO^ immortalized human lung fibroblasts (iHLFs). More than 25 cells from three independent experiments were analyzed for each condition (One-way ANOVA with post-hoc Sidak’s multiple comparison test). (C) Ratiometric quantification of active β3: active β1 integrin levels as determined by staining with MABT27 and 9EG7 antibodies, respectively. Images were masked using Paxillin counter-staining to restrict analysis to integrin adhesion complex (IAC) sites only. Data were collected from three independent studies (One-way ANOVA with post-hoc Sidak’s multiple comparison test). (D) Quantification of integrin adhesion complex (IAC) size as measured by total area of individual Paxillin puncta. Data were collected from three independent studies (Kruskal-Wallis test with post-hoc Dunn’s multiple comparison test). Data are represented as box-and-whisker plots of 10^th^-90^th^ percentiles with outlier points shown. For all statistical tests: ns = p > 0.05; p < 0.05 (*); p < 0.01 (**); p < .001 (***); p < .0001 (****).

### Thy-1 loss abrogates stiffness-dependent chromatin remodeling within promoter regions of mechanotransduction loci

While it has been reported that the Thy-1 locus is silenced by epigenetic modifications in idiopathic pulmonary fibrosis (IPF), the downstream changes in epigenetic activity and gene expression that contribute to persistent fibroblast phenotypic changes and disease progression following Thy-1 loss remain unknown.^31^ To characterize the divergences in fibroblast genomic activity after Thy-1^KO^ we executed a parallel workflow combining RNA-seq and ATAC-seq to capture dynamic changes in global gene expression and chromatin accessibility, respectively, exploring both stiffness- and time-dependent effects. Cells were cultured on Fn-coated tissue culture plastic (TCP; 1GPa) or hydrogels (3±1kPa and 22±2kPa) for periods of 24 and 72 hours. Technical replicates were collected in quadruplicate for each condition. All sequencing libraries passed appropriate quality control steps and were included in downstream analysis.

We first catalogued changes in the chromatin accessibility landscape that emerge after Thy-1 knockout using our ATAC-seq data. Unsupervised learning and multidimensionality reduction via Principal Component Analysis (PCA) revealed a clear segregation and clustering of samples based on both Thy-1 expression status (PC1; X-axis) and substrate stiffness (PC2; Y-axis) (Figure S2). Evaluation of extended PCs failed to reveal any additional insights or data trends. We then incorporated differential expression analyses into our ATAC-seq workflow to evaluate significant changes in chromatin accessibility within the Thy-1^KO^ population. This produced a comprehensive list of differentially accessible genomic loci (DALs) with significant changes between our two cell populations across stiffness and time. For all comparisons, >6,500 significant DALs (*p* ≤ 0.001) were found to exist within 5kb of a known transcription start site (i.e. within gene promoter regions). Promoter-associated DALs were then tested for gene enrichment based on their associated targets (Figure 2A). Interestingly, DALs were selectively and unilaterally enriched for coding loci associated with mechanical/integrin signaling pathways and regulation of cytoskeletal behavior. Directional examination (i.e. considering only upregulated or downregulated DALs in Thy-1^KO^ samples) found significant, bidirectional enrichment for ontologies tied to the actin cytoskeleton and FA componentry. These findings suggest that the aberrant cell spreading response seen in Thy-1^KO^ fibroblasts may stem from upstream changes in chromatin architecture that limit the capacity of these subpopulations to appropriately express the machinery necessary to spread in response to ECM mechanics.

**Figure 2.**
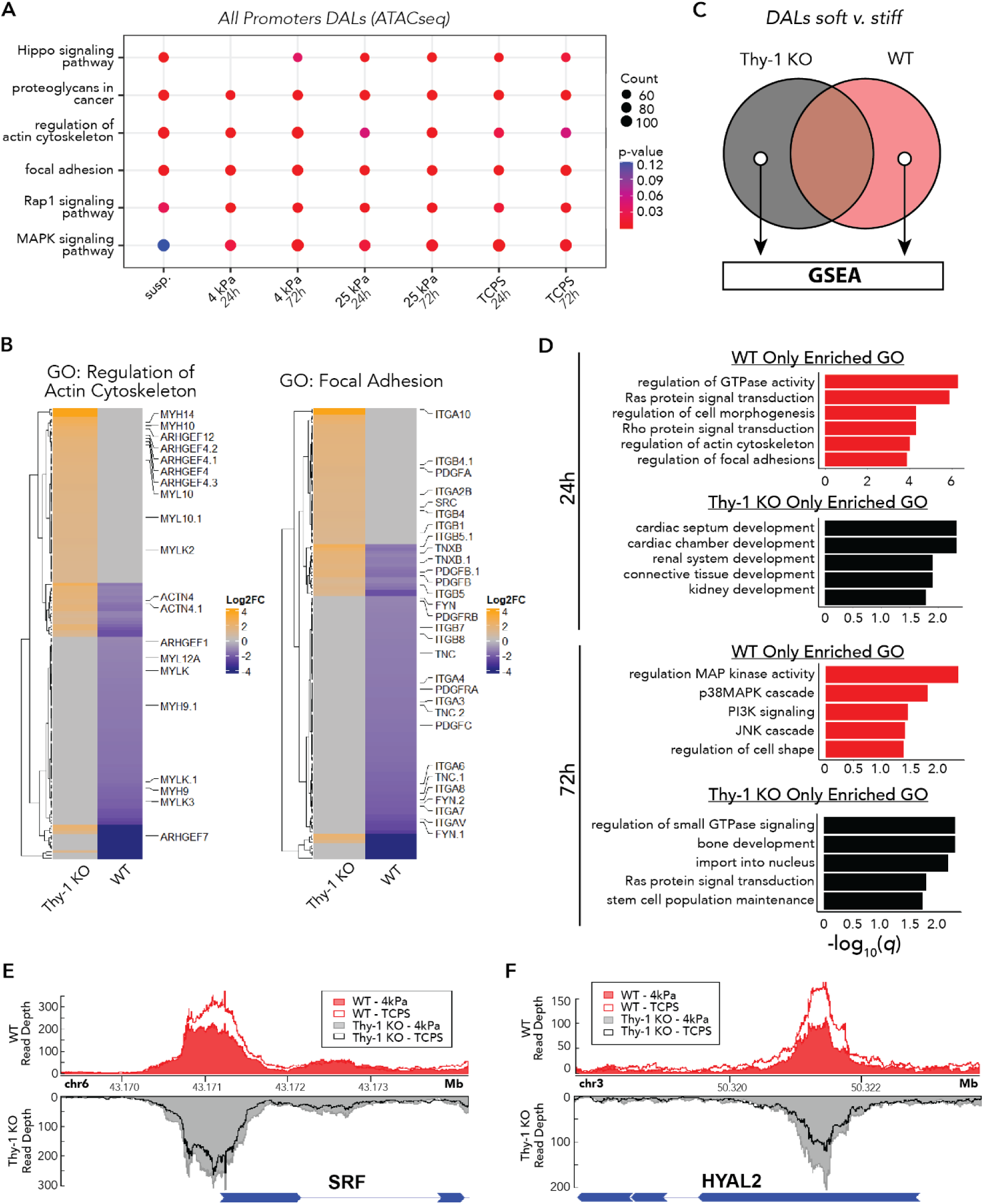
Impaired mechanoperception after Thy-1 loss induces aberrant patterns of chromatin remodeling at mechanotransduction promoter loci. (A) Kyoto Encyclopedia of Genes and Genomes (KEGG) enrichment analysis of differentially accessible loci (DAL) filtered to only include ATAC-seq peaks within 5kb of known transcription start sites. *q* < 0.05 was used to define significant enrichment (B) Heatmap of promoter accessibility changes between WT and Thy-1^KO^ lung fibroblasts for all members of the *Regulation of Actin Cytoskeleton* (left) and *Focal Adhesion* (right) ontologies (shading corresponds to Log2Foldchange of ATAC-seq read pileup for Thy-1^KO^/WT) Gene symbols with decimal points (e.g. MYLK.1) represent additional DALs identified within the same promoter region. (C) Schematic representation for identification of stiffness-dependent DALs from ATAC-seq data exclusive to each cell population. Promoter DALs between 4kPa and TCP were identified for each cell line at a given timepoint. The set difference between the two cell lines was then passed forward for enrichment analysis. (D) Gene ontology (GO) enrichment analysis for stiffness-dependent DALs exclusive to WT or Thy-1^KO^ iHLFs after 24 or 72hr in culture. *q* < 0.05 was used to define significant enrichment. Visualization of normalized ATAC-seq read pileup in WT (red) and Thy-1^KO^ (black/grey) fibroblasts at the (E) *SRF* and (F) *HYAL2* promoter regions after 24hr in culture (WT-4kPa = solid red fill; WT-TCP = red border transparent fill; Thy-1^KO^-4kPa = solid grey fill; Thy-1^KO^-TCP = black border transparent fill).

To better assess the promoters that contribute to FA and cytoskeletal organization enrichment in each cell line, we plotted the union set of DALs for each ontology after 72hr of culture on 4kPa hydrogels (Figure 2B). With respect to organization of the actin cytoskeleton, we identified 55 and 66 unique DALs in Thy-1^KO^ and WT fibroblasts, respectively. Thy-1^KO^ fibroblasts exhibited significantly upregulated promoter accessibility among members of the myosin heavy chain (*MYH*) family of motor proteins that generate contractile force in the cytoplasm. We also noted an enrichment among various *ARHGEF* species which are responsible for catalyzing the activation of Ras/Rho family members which orchestrate stress fiber formation and cellular contractility.^32^ Interrogation of the FA ontology revealed 42 and 78 DALs in Thy-1^KO^ and WT fibroblasts, respectively. Most critically, we noted the emergence of *FYN* and *SRC* as being differentially regulated between our two populations and is in agreement with our previous molecular observations showing increased Src recruitment and decreased Fyn recruitment to maturing adhesions after Thy-1 loss.^15^ Collectively, WT and Thy-1^KO^ fibroblasts utilize distinct modules of FA and cytoskeletal organization components to direct their spreading behavior via differential engagement of various SRC family kinase (SFK) members and GTPases that regulate cytoskeletal integrity.

While WT fibroblasts dynamically spread and accumulate myofibroblast activation markers as substrate stiffness increases, Thy-1^KO^ cells remain stagnant, displaying no such changes between soft and stiff culture conditions. As such, we aimed to evaluate whether these trends in mechanotransduction and cell spreading behavior were recapitulated in the chromatin organization of the Thy-1^KO^ genome. To this end we identified lists of promoter DALs whose accessibility were significantly changed between 4kPa and 1GPa substrate conditions in WT, but not Thy-1^KO^, and vice-versa for each timepoint (i.e. the set difference of WT/Thy-1^KO^ DALs and set difference Thy-1^KO^/WT DALs when comparing 4kPa vs. 1GPa; Figure 2C). When we considered DALs with significant, stiffness-dependent changes in promoter site accessibility at 24h in WT fibroblasts only we captured a strong enrichment towards mechanotransduction ontologies including Ras/Rho signaling, FA assembly, and actin cytoskeleton reorganization (Figure 2D). Thus, WT fibroblasts dynamically remodel promoter accessibility at mechanotransduction loci in response to substrate mechanics while Thy-1^KO^ retain similar accessibility profiles at these loci regardless of stiffness. After 72h in culture, a time-sensitive change in promoter activity was captured. In WT fibroblasts, FA and actin cytoskeleton-associated gene sets gave way to ontologies ascribed to MAPK pathway activation with a strong emphasis on stress-activated p38/JNK MAPK signaling that serve as effectors of TGF-β signaling.^33^ Interestingly, promoter DALs exclusive to Thy-1^KO^ samples at longer culture times did show enrichment for Ras/Rho signaling and nuclear import, which may potentiate the activity of known mechanoresponsive and myofibroblast-driving transcription factors such as MRTF, YAP/TAZ, and β-Catenin.^34, 35^

At the gene level, two distinct patterns of chromatin accessibility were noted for transcripts previously implicated in fibrotic progression. Genes such as *SRF*, a key MRTF co-factor that is requisite for the induction of pulmonary fibrosis in mouse models, displayed elevated levels of promoter accessibility in Thy-1 KO fibroblasts on soft substrates relative to WT but failed to respond to increases in substrate stiffness.^36^ By contrast, WT fibroblasts exhibited a dynamic, stiffness-dependent increase in *SRF* promoter accessibility, to a level higher than that of Thy-1 KO fibroblasts regardless of substrate stiffness (Figure 2E). A similar non-responsive Thy-1 KO accessibility trend was observed in the promoter of *CBX5* which acts as an epigenetic repressor recently shown to be essential in the maintenance of the myofibroblast phenotype (Figure S3A).^37^ We also captured a set of loci whose accessibility trends were inverted between WT and Thy-1 KO samples. *HYAL2,* which encodes a pro-myofibroblastic hyaluronidase, showed a positive correlation between substrate stiffness and *HYAL2* promoter accessibility in WT samples whereas Thy-1 KO displayed a negative correlation (Figure 2F).^38^ Only non-responsive trends (*MAP4K1/*MEKKK1 and *MAP2K1*/MEK1) in promoter accessibility were identified at 72hr timepoints (Figure S3B,C). Together, stiffness-dependent activation of promoter sites responsible for fibroblast activation in this dataset are exclusive to the WT genome during early stages of adaptation to the surrounding mechanical environment which later give way to biochemical signaling cascades responsible for carrying out normal mechanotransduction.

### RNA expression of genes tied to respiratory development and pattern specification are significantly downregulated following Thy-1 knockout

We next shifted our attention to profiling the transcriptomic response to Thy-1 loss in the fibroblast genome. Again, PCA revealed a clear segregation and clustering of samples based on both substrate stiffness (PC1; X-axis) and Thy-1 expression status (PC2; Y-axis) (Figure 3A). Unlike our ATAC-seq data, examination of extended PCs also revealed time-dependent clustering along PC3 (Figure S4). After 24h in culture, analysis of differentially expressed genes (DEGs; *q* < 0.05) across all substrate conditions identified at least 1,100 DEGs in each of the four comparisons between WT and Thy-1^KO^ samples (Range = [1,106, 2,227]; Figure 3B). Strikingly, a large cohort of DEGs (506 genes) were conserved independent of stiffness, suggesting a subset of dysregulated genes are driven solely as a function of Thy-1 loss. Similar trends were observed at the 72-hour timepoint and when comparing DEGs across various stiffnesses and timepoints (Figure S5). Among the set of 506 conserved DEGs, we noted a significant enrichment for mesenchymal development and embryonic patterning processes (Figure 3C). These findings collectively suggest that loss of Thy-1, regardless of substrate conditions, promotes widespread dysregulation of developmental processes tied to spatiotemporal regulation of organogenesis.

**Figure 3.**
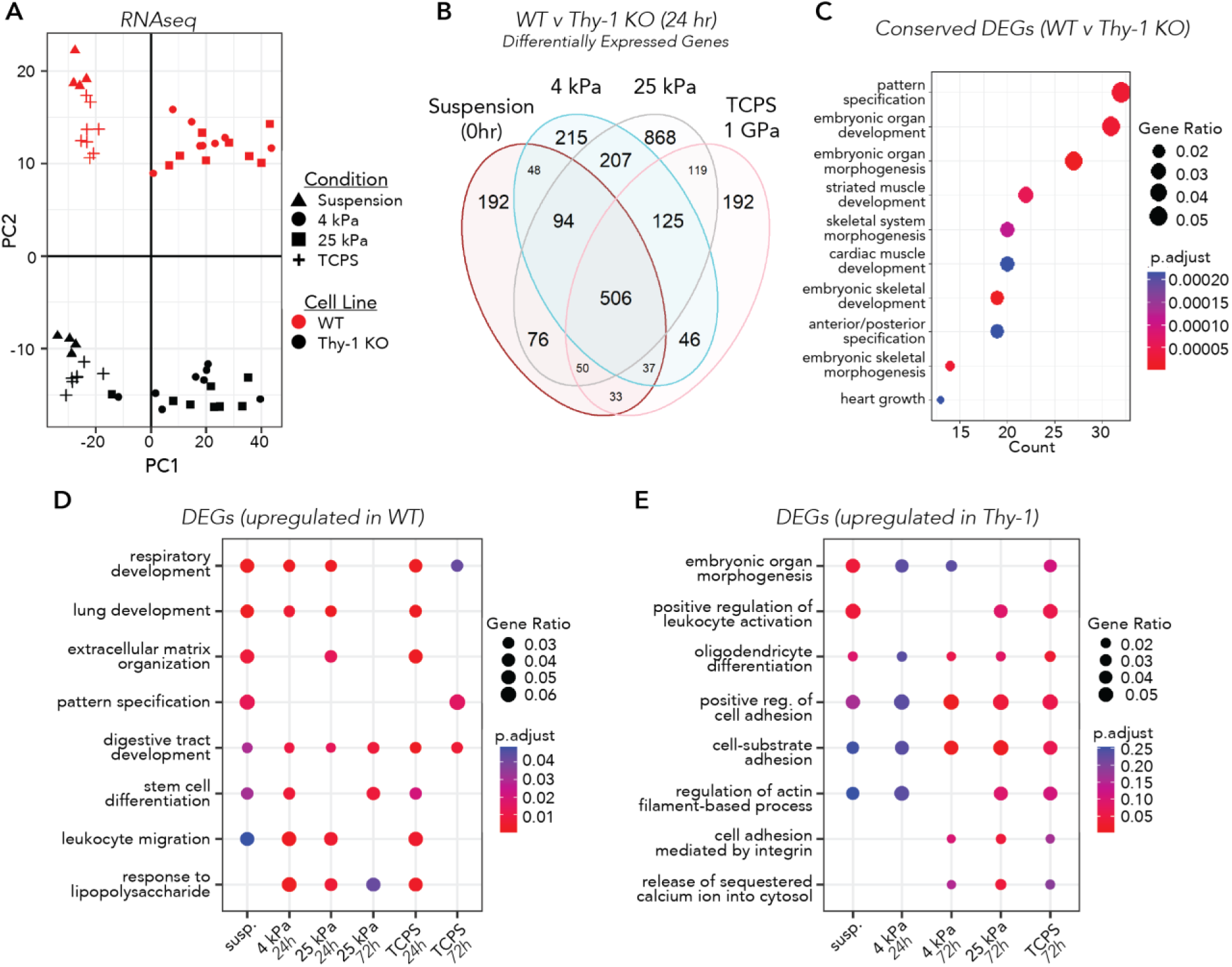
Impaired mechanoperception after Thy-1 loss leads to suppression of transcripts associated with embryonic patterning and pulmonary development. (A) Principal component analysis (PCA) of variance stabilized expression data based on weighted gene lists from RNA-seq output. X and Y-axes represent the primary and secondary principal components (PC1, PC2), respectively. PC1 = 51.58% of variance and PC2= 16.05% of variance. (B) Venn Diagram of differentially expressed genes (DEGs) between WT and Thy-1^KO^ immortalized human lung fibroblast cell lines (iHLF) for all substrate stiffnesses at 24hr (Wald test with BH correction; p < 0.05). A subset of 506 DEGs were conserved across all comparisons while a range of DEGs [192, 868] were uniquely expressed in a substrate dependent manner. (C) Gene ontology (GO) enrichment analysis of 506 conserved DEGs after Thy-1 loss at 24hr. (D,E) GO enrichment analysis for DEGs upregulated in (D) WT or (E) Thy-1^KO^ fibroblasts across all displayed comparisons (comparisons not listed exhibited no significant changes in GO-BP pathways). Adjusted *p* < 0.05 was used to define significant enrichment

Condition-wide gene set enrichment analysis was performed on our DEG lists to connect trends in gene expression to changes in cell function and behavior as a function of substrate stiffness and culture time. Among DEGs upregulated in WT fibroblasts, again, there was near unilateral enrichment for various developmental processes. Strikingly, we noticed the emergence of new terms with previous relevance to Thy-1 biology including stem cell differentiation and lung development (Figure 3D). In contrast, upregulated DEGs in Thy-1^KO^ displayed no concordant enrichment across time or stiffness (Figure 3E). After 72h in culture, Thy-1^KO^ fibroblasts displayed an upregulation of processes tied to cell adhesion, actin cytoskeleton organization, and integrin signaling. These data shed additional light upon prior findings related to the impact of Thy-1 knockout on fetal lung development, here, illustrating that Thy-1 loss in pulmonary fibroblasts promotes a significant decrease in transcription of genes necessary for proper maturation of the developing lung.

### Silencing of the HOXA family of tissue patterning transcription factors is the distinguishing feature of the Thy-1^KO^ genome

Quantification of gene-by-gene loadings along PC2 of our RNA-seq PCA (i.e. the PC that clustered samples based on Thy-1 expression status) identified *HOXA5 and HOXA3* among the top 10 values contributing strongest to cluster segregation (Figure 4A). Interrogation of the top 50 loading values along PC2 revealed additional *HOXA* species (*HOXA7, HOXA2, HOXA4,* and *HOXA1*; Table S1). Interestingly, each of these genes were downregulated in Thy-1^KO^ samples, suggesting a direct link between loss of mechanoperception and suppression of *HOXA* cluster expression. Visual examination of the trends in gene expression (Figure 4B, bottom panel), log_2_(Thy-1^KO^/WT) foldchange (Figure 4B, blue-red heatmap), and statistical significance of differential gene expression (Figure 4B, yellow-brown heatmap) of all *HOXA* mRNA species revealed *HOXA5* as both the most significantly downregulated and most highly expressed member of the *HOXA* cluster. It is also worth noting that there was a gradual shift in *HOXA* regulation between the anterior (*HOXA1-6)* and posterior (*HOXA7-13*) genes. This may be associated with the 3-dimensional organization of the genome as a topology associated domain (TAD) has been reported to exist between the anterior and posterior *HOXA* genes in lung fibroblasts and serves to decouple their regulation.^39^

**Figure 4.**
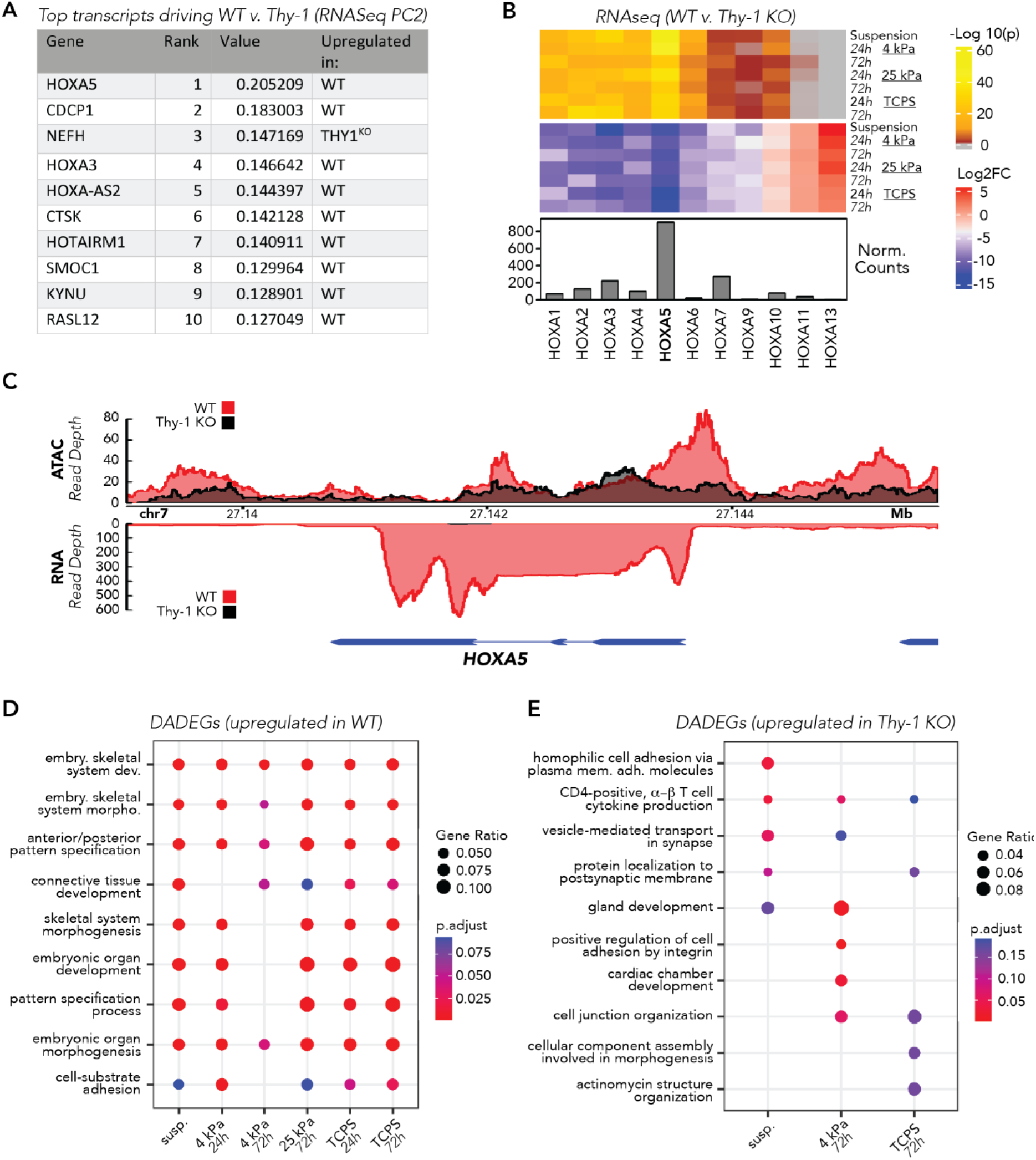
Silencing of the *HOXA* family of developmental factors is a defining genomic signature of the Thy-1^KO^ pulmonary fibroblast. (A) Gene-wise rank of loading values contributing to RNA-seq PC2 which separates samples on Thy-1 expression status. (B) Heatmap of Benjamini-Hochberg corrected p-values (top; grey = ns) and log2 foldchange (bottom) for protein coding *HOXA* family members. Bottom annotation represents DESeq2 normalized transcript counts for each gene. Foldchanges are reported as log2(Thy-1^KO^/WT). (C) Representative plot of ATAC-seq (top) and RNA-seq (bottom) read pileup at the *HOXA5* locus for WT (red) and Thy-1^KO^ (black) iHLFs cultured on 4kPa hydrogels for 24hr. (D,E) GO-BP analysis of loci showing concurrent upregulation of gene expression and *cis-*regulatory accessibility (differentially accessible, differentially expressed genes; DADEGs) in WT samples and Thy-1^KO^ samples, respectively. Adjusted *p* < 0.05 was used to define significant enrichment.

We next evaluated our ATAC-and RNA-seq datasets in combination to validate our observations of *HOXA5* expression patterns and gain further insights into how changes in chromatin accessibility may regulate corresponding changes in gene expression. As chromatin accessibility is diminished via the condensation of DNA around histone octamers, steric hindrance physically restricts transcription factors (TFs) from binding DNA and regulating gene activity.^40^ Thus, in the case of promoter and enhancer sites, loss of chromatin accessibility is correlated with a reduction in gene expression. We overlayed pileup of ATAC and RNA sequencing reads along the *HOXA5* locus after culture on 4kPa hydrogels for 24h in both WT and Thy-1^KO^ fibroblasts (Figure 4C). In agreement with our PCA loadings, we identified a stark drop-off in both chromatin accessibility (top) and RNA transcripts (bottom) in the Thy-1^KO^ condition. Similar trends held, to varying degrees, for all other comparisons across stiffness and time (Figure S6). No such trends were observed at the *HOXB5* or *HOXC5* loci, implying that Thy-1 loss selectively targets *HOXA5* and compensation of other *HOX5* species does not ensue (Figure S7). These data suggest that Thy-1 loss alone was sufficient to stimulate a drastic shift in epigenetic activity at the *HOXA5* locus that inhibits TF access to the gene’s promoter site and induces a near complete loss of transcriptional activity as a result.

To evaluate whether our observations at the *HOXA5* locus were concentrated to a single developmental factor or indicative of a larger shift in cell behavior we utilized the *BETA* package to connect significant changes in gene expression (RNA-seq) with concordant changes in chromatin accessibility at *cis*-regulatory loci (hereafter termed: differentially accessible differentially expressed genes; DADEGs).^41^ *BETA* analysis was applied to generate tables of significant (rank product ≤ 1e-3) DADEGs for all comparisons and enrichment analyses were performed on the resulting lists (Figure 4D, E). When considering DADEGs downregulated in Thy-1^KO^ fibroblasts, we noted a significant enrichment towards embryonic development and morphogenic processes that was retained across all comparisons. More interestingly, there was a highly significant enrichment in pattern specification processes which are directly linked to HOX activity. Evaluation of DADEGs upregulated in Thy-1^KO^ fibroblasts failed to reveal any conserved enrichment trends. Culture on 4kPa substrates for 72hr exhibited enrichment towards integrin signaling and cell adhesion processes, which aligns well with our prior discovery that Thy-1 loss led to enhanced promoter site accessibility at loci tied to Ras/Rho signaling and cytoskeletal tension relative to WT fibroblasts in soft substrate conditions. In sum, *HOXA5* promoter and transcript silencing appears to be downstream of Thy-1 knockout and may induce a fundamental shift in fibroblast genome activity away from developmental and regionalization processes which could have profound implications on the mechanisms by which Thy-1^KO^ human lung fibroblasts respond to wound healing cues in the lung.

### Identification of HOXA5 as a mechanoresponsive transcription factor in pulmonary fibroblasts

We next aimed to validate and characterize the patterns of HOXA5 protein levels as a function of substrate rigidity. Plotting of normalized transcript counts in WT fibroblasts after 24h in culture showed a significant loss in *HOXA5* expression on 1GPa substrates but no differences between 4kPa and 25kPa hydrogels (Figure 5A). These data indicate that either Thy-1 loss or increasing substrate stiffness, both of which are known drivers of myofibroblast activation, are negative regulators of *HOXA5* gene activity. To confirm that loss of *HOXA5* on 1GPa substrates was indeed driven by stiffness, we assessed HOXA5 protein levels on 5kPa and 100kPa hydrogels in addition to 1GPa TCP. Substitution of 100kPa vs. 25kPa hydrogels provides a more pronounced stiffness response while still controlling for substrate composition. Indeed, in WT fibroblasts, HOXA5 showed a significant reduction in signal as substrate stiffness increased (Figure 5B, C). Thy-1^KO^ fibroblasts, regardless of culture substrate stiffness, expressed HOXA5 protein at levels similar to or below WT fibroblasts cultured on the highest culture substrate stiffness (TCPS, 1GPa). Similar results were observed for multiple Thy-1^KO^ clones, illustrating that this is not an artifact of CRISPR-Cas9 transfection nor of clonal expansion during Thy-1^KO^ cell line generation (Figure S8). These protein level findings corroborate our sequencing observations and further capture an inverse correlation between substrate stiffness and HOXA5 protein levels in WT lung fibroblasts. The latter is of particular interest given that Thy-1 loss induces a loss of mechanoperception and induces cell spreading phenotypes akin to WT fibroblasts cultured on stiff ECM. This suggests that cytoskeletal organization may be a key determinant of HOXA5 expression. We tested this hypothesis by treating WT and Thy-1^KO^ fibroblasts seeded on TCPS (1GPa) substrates with Latrunculin A (LatA), which catalyzes actin depolymerization, or Rho-associated kinase (ROCK) inhibitor Y-27632, which reduces the contractility of cells. Both LatA and Y27632 induced an upregulation of HOXA5 expression (Figure 5D, E), with LatA having a more profound effect on WT and Y27632 having a greater impact on Thy-1^KO^. Given that RhoGEF promoter site accessibility was a defining feature in Thy-1^KO^ fibroblasts, these incongruent observations may be due to deviations in promoter usage at loci associated with FA formation and actin cytoskeleton regulation previously noted from our ATAC-seq analysis (Figure 2).

**Figure 5.**
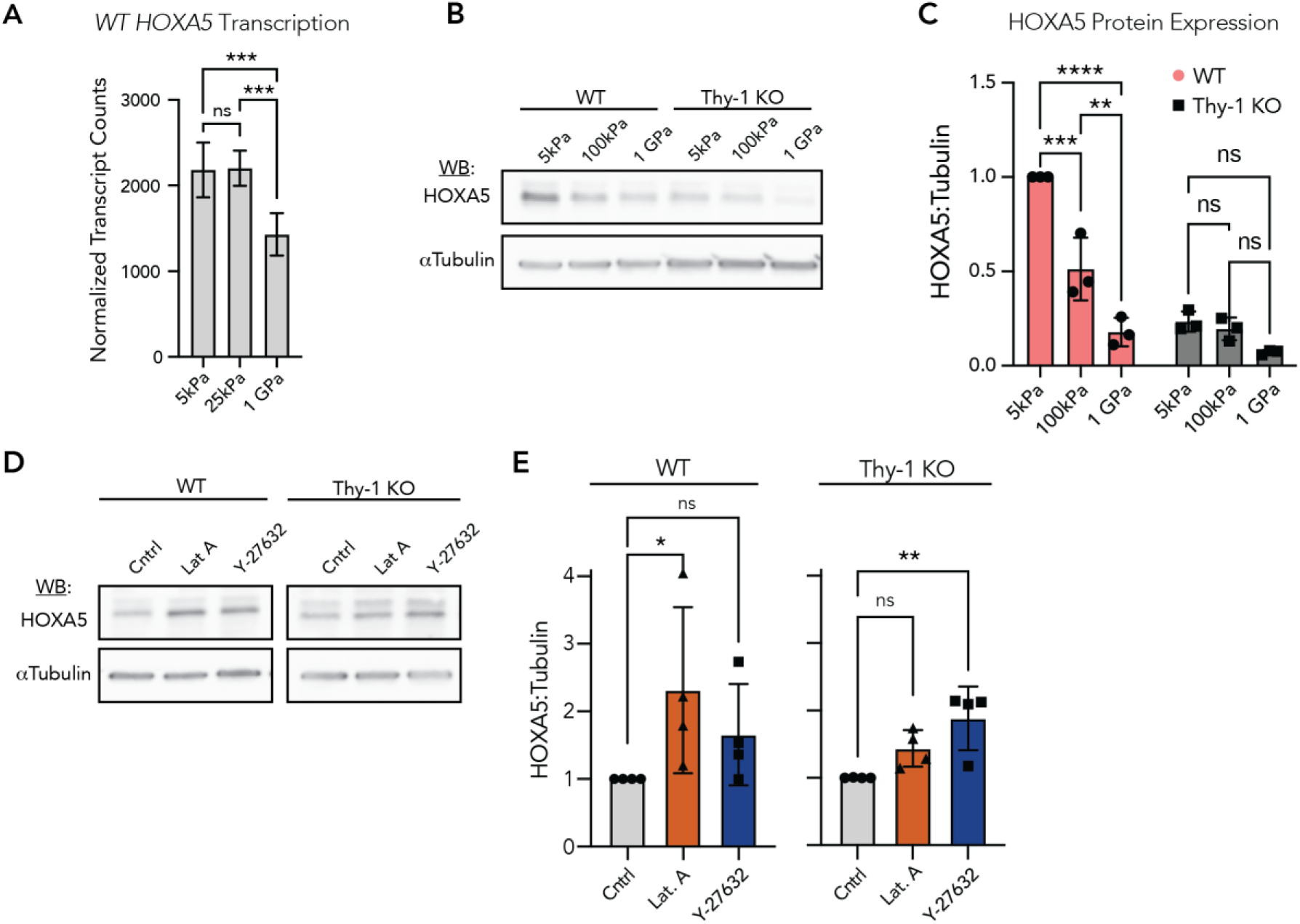
HOXA5 expression is regulated by perceived substrate stiffness and cytoskeletal integrity. (A) DESeq2 normalized transcript counts in WT iHLFs for each stiffness after 24hr in culture. Mean ± S.D. plotted; N=4 technical replicates; 1-way ANOVA with post-hoc Tukey’s multiple comparison test. (B) Western blot for HOXA5 in WT and Thy-1^KO^ iHLFs cultured on 5kPa and 100kPa PDMS hydrogels as well as TCP (∼1GPa). (C) Quantification of HOXA5 normalized to loading control and plotted as a ratio of normalized signal relative to WT-5kPa (Mean ± S.D. plotted; N=3 independent experiments; 2-way ANOVA with post-hoc Tukey’s multiple comparison test). Values are reported as normalized HOXA5 signal relative to WT-5kPa within each technical replicate. (D) Western blots for HOXA5 in WT and Thy-1^KO^ iHLFs cultured on TCP following stimulation with 1μM actin polymerization inhibitor Latrunculin A or 10μM ROCK inhibitor Y-27632 for 3 and 6hr, respectively. (E) Quantification of HOXA5 normalized to loading control and plotted as a ratio of normalized signal relative to DMSO vehicle control (Mean ± S.D. plotted; N=4 independent experiments; non-parametric Kruskal-Wallis test with post-hoc uncorrected Dunn’s test) ns = *p* > 0.05; *p* < 0.05 (*); *p* < 0.01 (**); *p* < .001 (***); *p* < .0001 (****).

### SRC family kinase (SFK) signaling and integrin α_v_ promoting environments mediate HOXA5 expression in a substrate dependent manner

Having established a link between HOXA5 protein levels and substrate stiffness, we next sought to identify FA signaling mechanisms that might explain, in part, HOXA5 downregulation on stiff substrates. Previous research has shown that loss of Thy-1 results in increased integrin α_v_ affinity for ECM and a concurrent increase in recruitment of c-Src to FAs.^15^ Thus, we first evaluated whether c-Src and the broader SRC family of kinases (SFKs) are involved in the control of HOXA5 protein expression. Treatment with the c-Src specific inhibitor KB SRC 4 or the pan-SFK inhibitor PP2 were both sufficient to induce a significant increase in HOXA5 expression on stiff, myofibroblast promoting ECM in both cell lines (Figure 6A). Neither inhibitor had a statistically significant effect on either cell line on 5kPa hydrogels, though HOXA5 expression in Thy-1^KO^ cells approached statistical significance (*p* = 0.06). For all conditions, similar trends were observed after treatment with inhibitors for 3h (Figure S9). The complete lack of a response to c-Src and SFK inhibitor treatment in WT fibroblasts on soft substrates corroborates prior literature demonstrating extremely low baseline levels of SFK activity in weakly spread cells.^15^ This result further supports the hypothesis that activation of c-Src and perhaps additional SFKs are involved in the inhibition of HOXA5 expression on stiff ECM.

**Figure 6.**
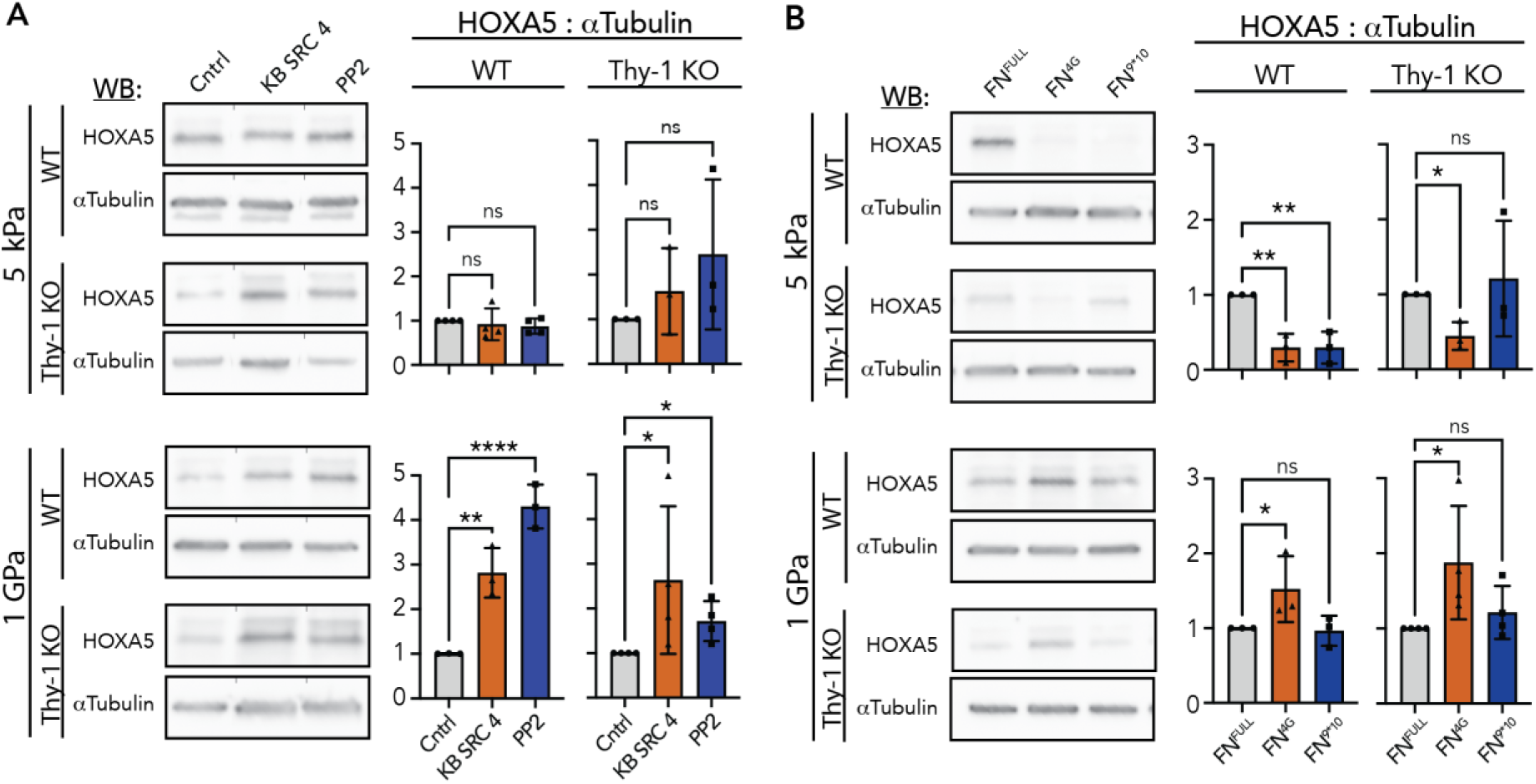
Src family kinase activity and fibronectin (Fn)-binding integrin signaling are upstream regulators of HOXA5 expression dynamics. (A) Western blots for HOXA5 in WT and Thy-1^KO^ iHLFs cultured on 5kPa PDMS hydrogel or TCP following stimulation with 20μM SRC specific inhibitor KB SRC 4 or 10μM pan-SFK inhibitor PP2 for 6hr. Quantification of HOXA5 normalized to loading control was plotted as a ratio of normalized signal relative to DMSO vehicle control. Mean ± S.D. plotted; N=3 (Thy-1^KO^-5kPa and WT-1GPa) or N=4 (WT-5kPa and Thy-1^KO^-1GPa) independent experiments; WT-5kPa and WT-1GPa = 1-way ANOVA with post-hoc uncorrected Fisher’s LSD test, Thy-1^KO^ -5kPa and Thy-1^KO^-1GPa = non-parametric Kruskal-Wallis test with post-hoc uncorrected Dunn’s test. Tests chosen based on data distribution and variance. (B) Western blots for HOXA5 in WT and Thy-1^KO^ iHLFs cultured on 5kPa PDMS hydrogel or TCP substrates coated with αV-promoting (Fn^4G^) or α5-promoting (Fn^9*10^) peptide fragments of the 9-10FnIII integrin binding site. Quantification of HOXA5 normalized to loading control was plotted as a ratio of normalized signal relative to full length human Fn (Fn^Full^) control. Mean ± S.D. plotted; N=3 (WT-5kPa, WT-1GPa, and Thy-1^KO^-5kPa) or N=4 (Thy-1^KO^-1GPa) independent experiments; WT-5kPa = 1-way ANOVA with *post-hoc* uncorrected Fisher’s LSD test, Thy-1^KO^ -5kPa = Brown-Forsythe and Welch ANOVA with *post-hoc* unpaired t-tests with Welch’s correction, WT-1GPa and Thy-1^KO^ -1GPa = non-parametric Kruskal-Wallis test with *post-hoc* uncorrected Dunn’s test. Tests chosen based on data distribution and variance. For all statistical tests: ns = *p* > 0.05; *p* < 0.05 (*); *p* < 0.01 (**); *p* < .001 (***); *p* < .0001 (****).

Based on prior descriptions of Thy-1’s role in the regulation of integrin α_v_-mediated adhesion and downstream hyperactivation of c-Src, we next sought to test whether upregulated adhesion through integrin α_v_ on soft substrates is sufficient to inhibit HOXA5. To achieve this, we employed the use of engineered fragments of Fn’s integrin binding domain, specifically the 9^th^ and 10^th^ type III repeats of Fn.^42^ Briefly, Fn^9*10^ presents a previously reported point mutation that structurally stabilizes the native conformation of the integrin binding domain and promotes the native cell-Fn interaction with integrin α_5_β_1_ while maintaining a weaker affinity for integrin α_v_β_3_. In contrast, Fn^4G^ possesses a flexible linker region that disrupts Fn binding to α_5_β_1_ while affinity to integrin α_v_β_3_ remains unaffected. HOXA5 protein expression in fibroblasts cultured on 5kPa substrates presenting the α_v_β_3_-favoring Fn^4G^ demonstrated a robust and significant loss in HOXA5 expression compared with full length Fn controls for both cell lines (Fn^Full^, Figure 6B). Surprisingly, adhesion of fibroblasts to Fn^9*10^ was sufficient to retain native HOXA5 expression levels in Thy-1^KO^ fibroblasts, but not in WT fibroblasts where expression was significantly reduced. No changes in HOXA5 levels were observed on either Fn protein fragment when cells were cultured on 100kPa hydrogels (Figure S10). On 1GPa TCPS, Fn^4G^ was found to promote an increase in HOXA5 expression in both cell types. Notably, both fibroblast populations display a more poorly spread phenotype on TCPS presenting Fn^4G^ relative to other Fn species, further suggesting the necessity of cytoskeletal activation/signaling in stiffness-mediated HOXA5 inhibition. Collectively, these findings show that disruption of Fn-binding integrin stoichiometry is sufficient to impair stiffness-dependent HOXA5 expression dynamics.

## Discussion

Cell-ECM interactions, adhesion complex dynamics and mechanoperception/ mechanotransduction are highly relevant in human health and disease, particularly in scenarios of tissue formation during development and tissue regeneration/remodeling responses after wounding. Our study helps address this gap by identifying a potential link between fibroblast mechanotransduction and a genomic programming switch that balances pro-regeneration/tissue homeostasis and pro-fibrotic/tissue destruction phenotypes. Based on our data, this programming switch can be subverted by disrupting mechanisms of normal fibroblast mechanoperception (e.g. loss of Thy-1 expression). Impaired mechanoperception directly impacts chromosomal structure, effectively hard-wiring an aberrant phenotype by restructuring the fundamental make up of cell-ECM interactions and thus eliminating phenotypic plasticity linked to microenvironmental mechanical signals.

In this manuscript, we compared WT and Thy-1^KO^ fibroblasts, as one of potentially many examples of cellular mechanoperception dysfunction. Insights on changes in chromatin architecture from our ATAC-seq dataset revealed dysregulated promoter-site accessibility at loci connected to FA machinery and remodeling of the actin cytoskeleton following Thy-1 knockout. In Thy-1^KO^ fibroblasts cultured on soft substrates, we noted elevated promoter accessibility at various *ARHGEF* family members that catalyze the activation of Rho family GTPases responsible for lamellipodial protrusion and F-actin stress fiber formation.^43^ Thus, upon Thy-1 loss, fibroblasts are epigenetically primed to acquire a well-spread, contractile phenotype, even in soft environments. An interesting, and confounding, aspect of the Thy-1^KO^ phenotype is its inability to further spread and respond normally to biophysical cues such as increased ECM stiffness. This may be explained by the fact that while WT fibroblasts display stiffness-dependent changes in promoter site accessibility at pro-myofibroblastic loci, Thy-1^KO^ fibroblasts do not. Stiffness-mediated myofibroblast differentiation is a defining feature of fibroblasts throughout the literature and these findings suggests that the defect in mechanoperception following Thy-1 loss traps the fibroblast in a static state of mechanical purgatory.

In addition to refining our understanding of the epigenetic contributions to the aberrant cytoskeletal phenotypes known to emerge in response to Thy-1 loss, we found that the most distinct and recurrent feature of the Thy-1^KO^ genome was a broad silencing of transcripts and regulatory sites important in organ system development as well as tissue and embryo patterning. The most prominent example was the systemic closure of promoters of the anterior *HOXA* family which, in adult tissues, are proposed to provide a spatial localization signature that contributes to the maintenance of tissue-specific phenotypes.^44–46^ An initial high-level interpretation might suggest Thy-1 augments pulmonary fibroblast maturation in the developing lung and helps maintain the fibroblast’s sense of regionalization in adult tissues. This is underscored in our data by near complete silencing of *HOXA5,* a pioneer factor that ascribes the spatial orientation of the thoracic space and is necessary for pulmonary development, following Thy-1 knockdown. Though a thorough interrogation of the genomic mechanism by which *HOXA5* is suppressed following Thy-1 loss is outside the scope of this work, there is strong evidence that increased activity of the polycomb repressive complex (PRC), a well described negative regulator of *HOX* gene expression, is implicated in our system. Imputation of *cis* binding factors tied to downregulated DEGs following Thy-1 loss revealed a number of PRC components (SUZ12, EZH1/2, JARID2) as critical regulators of gene repression in Thy-1 KO fibroblasts across all stiffnesses and timepoints (Table S2). A critical question that emerges from these findings is how HOXA5 silencing may impact fibroblast function in neonatal and adult tissues. Anecdotally, two separate *in vivo* studies evaluating *Hoxa5*^+/-^ and *Thy1*^-/-^ neonatal lung development and function characterized strikingly similar histological abnormalities including alveolar simplification and aberrant positioning of αSMA^+^ myofibroblasts at identical developmental timepoints.^24, 47^ Our analyses at the RNA and protein levels showed an inverse correlation between HOXA5 expression and conditions that promote myofibroblast activation, a relationship completely disrupted by Thy-1 loss. This is given further credence by our discovery of a global shift in chromatin organization that resulted in widespread closure of HOXA5 DNA binding sites follow Thy-1 KO on soft, but not stiff, ECM (Figure S11). Thus, it is plausible that Thy-1 loss-induced dysregulation of FA integrin stoichiometry and signaling is a mechanism that induces HOXA5 repression and a loss of lung-specific functions within lung fibroblasts which would have dire consequences in lung injury repair.

Soft substrates, which normally enable the expression of HOXA5 in WT fibroblasts, coated with an ECM ligand strongly favoring integrin α_v_ engagement and signaling (i.e. Fn^4G^) induced a marked reduction in HOXA5 protein expression. At a molecular level, this implicates the stoichiometry of Fn-binding integrins within FAs, rather than ECM mechanics alone, as essential to the maintenance of HOXA5. Yet, the near equivalent level of HOXA5 inhibition in fibroblasts engaging an engineered ECM ligand supporting the complete repertoire of Fn binding integrins (i.e. Fn^9*10^) suggests an additional signal from the FA is required for HOXA5 maintenance. When engaging the same integrin α_v_ promoting ligands coated on HOXA5-silencing stiff substrates, both WT and Thy-1^KO^ fibroblasts upregulated expression of HOXA5. Given that the Fn^4G^ engineered fragment of Fn does not support strong adhesion, FA maturation, or contractile force generation, this result corroborates the active role of FA maturation and cytoskeletal contractility in the HOXA5 regulatory axis.

Our study points to an emerging genomic circuit that links mechano- and ECM-responsive chromatin organization to the repression of pulmonary development transcripts in lung fibroblasts that clearly warrants further investigation. Carefully designed experiments outside the scope of the current work will be required to fully elucidate this novel mechanism linking ECM adhesion dynamics and mechanoperception to key factors defining a fibroblast’s tissue type, positioning, patterning, and perhaps stemness. Given the known connections between aging and fibrosis, alongside the abnormal developmental patterns observed in Thy-1 and HOXA5 mutant embryos, it is quite likely that a connection between fibroblast activation state and HOXA5 expression must be tightly controlled to carryout (re)generative processes critical to pulmonary tissue formation and maintenance. HOXA5’s reported role in restricting fibroblast contraction and promoting apoptosis could position adhesion-mediated regulation of HOXA5 as a critical breakpoint in the regeneration v. fibrogenesis genetic code.

## Materials/Methods

### Cell Culture, flow cytometry, and GPI-aIR gene knock out using CRISPR-CAS9

Immortalized human lung fibroblasts (iHLF) were commercially purchased (ABM®) and maintained in DMEM supplemented with 10% FBS, 100μg/ml penicillin/streptomycin, and 1mM sodium pyruvate (normal growth medium). Surface expression of Thy-1 was confirmed via flow cytometry with a FITC-conjugated mouse anti-human CD90 monoclonal antibody (1:100, FITC mouse anti-human CD90, clone 5E10, BD Biosciences). To generate Thy-1 knock-out (Thy-1^KO^) fibroblasts, iHLF were transfected with the Alt-R® CRISPR-CAS9 system (IDT; Coralville, IA) per the manufacturer’s instruction and then single cell sorted into a 96-well plate based on the tracRNA marker ATTO 550 using a FACSAria™ Fusion sorter (BD Biosciences; Lakes, NJ). Thy-1^KO^ was confirmed with T7EI kit (IDT) and loss of surface expression of Thy-1 was verified via flow cytometry (1:100, FITC mouse anti-human CD90, clone 5E10, BD Biosciences).

### Hydrogel preparation

Soft (2kPa) and stiff (20kPa) polyacrylamide (PAA) hydrogels were prepared following a previously published protocol.^15^ Briefly, 12 mm glass coverslips were plasma-etched in a plasma cleaner for 45s. Following the treatment, coverslips were coated with freshly made 5% APTES in absolute Ethanol for 10 min at room temperature. APTES was then removed, and coverslips were thoroughly washed with diH_2_O. After APTES treatment, coverslip surfaces were coated with a 0.5% Glutaraldehyde in H_2_O solution for 30 min at room temperature, followed with washing. Acrylamide/bis-acrylamide mixtures were prepared according to the corresponding stiffness recipe and de-gassed. For each coverslip, 15uL of PAA was mixed with .015uL TEMED and 0.45μL 10% APS then vortexed to initiate crosslinking. The PAA mixture was immediately pipetted onto a DCDMS-treated slide with the APTES/Glutaraldehyde-treated coverslip inverted and placed, treatment side down, onto the PAA mixture. Hydrogels were allowed to crosslink and cure for 1-2hr at room temperature. After gel formation, hydrogels were functionalized with Sulfo-SANPAH and coated with 10μg/mL full length human Fn at 37C overnight.

### Cell morphology and integrin ratio metric study

WT and Thy-1^KO^ cells were seeded on 2 or 20 kPa PAA hydrogel or glass control coverslips (∼2 × 10^4^ cells/cm2) coated with 10μg/mL full length human Fn and cultured at 37C for 2hr before fixation with 4% paraformaldehyde followed with permeabilization and overnight blocking. Cells were then stained with anti-Paxillin antibody (Y113, Abcam) or a combination of anti-active αvβ3 (MABT27, EMD Millipore) and anti-active α5β1 (9EG7, BD Biosciences) overnight followed by co-staining with respective secondary antibodies for 1hr at room temperature. Cells were then subjected to immuno-fluorescent microscopy with a Perkin-Elmer spinning disk Ultra-View system with a 60X, 1.49 N.A. oil immersion objective and quantified using Volocity (Perkin Elmer). Adhesion complexes from at least 50 cells from each condition were quantified and analyzed.

### RNA-seq and ATAC-seq library preparation

WT and Thy-1^KO^ cells were seeded on 4kPa polyacrylamide hydrogels (Matrigen), 25kPa polyacrylamide hydrogels (Matrigen), or tissue culture plastic (TCP) pre-coated with 10μg/mL full length human Fn. RNA and DNA were collected concurrently after culture periods of 24 or 72 hr. Prior to seeding, a baseline aliquot of cells was immediately collected for downstream library synthesis (labeled as “suspension” in downstream analyses). ATAC-seq libraries were generated from 100k cells following previously established protocols.^48^ Samples were double-barcoded using Nextera i7 and i5 indexes (Illumina), size selected to exclude fragments >600 bp using the SPRIselect system (Beckman-Coulter) and pooled to a final concentration of 20nM/library. Proper fragment size distribution was validated prior to sequencing via TapeStation (Agilent) as preliminary quality control (QC) to confirm DNA fragment sizes followed the anticipated pattern of nucleosomal periodicity. For RNA-seq, cells were lysed using TriZol (Thermo Fischer) and RNA isolated with Qiagen’s RNeasy Micro Prep kit. Isolated RNA was converted into sequencing compatible libraries using the TruSeq Stranded mRNA Library Prep kit (Illumina). RNA isolates with masses >= 0.5μg and RNA integrity number (RIN) scores >= 9.0 were sequenced. Both RNA-seq and pooled ATAC-seq libraries were sequenced to a depth of ∼20 M paired-end reads (2x150 bp) using the Illumina NovaSeq6000 platform.

### RNA-seq data processing and analysis

Raw fastq files were first adapter trimmed and passed for preliminary QC using the *TrimGalor*e! and *FastQC* packages, respectively.^49, 50^ Trimmed reads were then mapped to the hg38 genome using *HISAT2* and converted into gene count tables via *Stringtie* and *Ballgown* as previously described.^51^ All processed samples passed preliminary QC and were passed into downstream analyses using R. Differential gene expression analysis was performed between conditions using *DESeq2* with a BH-adjusted p-value < 0.05 to denote genes with significant expression changes between conditions.^52^ Principal component analysis (PCA) was quantified from variance stabilized *DESeq2* results. Heatmaps of gene expression were generated using the *ComplexHeatmap* package from *DESeq2*-normalized read counts after filtering out genes whose mean expression amongst all samples totaled < 10 reads.^53^ Venn Diagrams of overlap amongst differentially expressed genes were generated and visualized using *Limma*.^54^ All enrichment analyses and dot plot visualizations were performed using the clusterProfiler package with a significant *q-*value threshold of 0.1.^55^ The Binding Analysis for Regulation of Transcription (*BART*) package was applied to impute potential transcriptional regulators of DEGs identified from *DESeq2* analyses.^56^ Regulators with Irwin-Hall p-value < 0.01 were considered significant.

### ATAC-seq data processing and analysis

As with RNA-seq, raw fastq files were first adapter trimmed and passed for preliminary QC using *TrimGalore*! and *FastQC*. Trimmed reads were mapped to the hg38 genome using the *Bowtie2* aligner.^57^ Peaks were called on aligned reads using the --call-peaks function from the *MACS2* package and peak summits identified as the region stratifying 100 bp upstream and downstream of each peak’s center (201 bp total).^58^ For all samples, peak summits identified using *MACS2* were merged into a singular bam file and converted into a count matrix of read accumulation within peak regions on a sample-by-sample basis using the package *featureCounts*.^59^ Principal component analysis (PCA) was performed using variance stabilized *DESeq2* results. For promoter enrichment analyses, *HOMER* was employed to quantify differential accessibility and the resulting loci were annotated using the *GenomicRanges* package.^60, 61^ Peaks displaying a p-value < 1e-3 between test conditions were filtered to restrict analysis to loci whose absolute distance from the nearest transcription start site was <= 5kb. Reads found within peaks that exhibited significant changes in binding accessibility between conditions were converted to fasta file format and passed to the *HOMER* package to evaluate transcription factor binding site (TFBS) enrichment analysis.^60^ All enrichment analyses and dot plot visualizations were performed using the *clusterProfiler* package with a significant *q-*value threshold of 0.1.^55^ Heatmaps of promoter enrichment data were generated using the *ComplexHeatmap* package.^53^ The *Sushi* package was applied to plot ATAC-seq traces along the hg38 genome.^62^ HOXA5 transcription factor binding site accessibility was quantified using *MEME* Suite.^63^ ATAC-seq peaks identified from MACS2 output with WT samples were used as inputs for the test condition and Thy-1^KO^ peaks as the background condition.

### Merged analysis of RNA-seq and ATAC-seq results using BETA

The Binding and Expression Target Analysis (*BETA*) package from the Cistrome project was incorporated to merge RNA-seq and ATAC-seq results.^41^ Briefly, *BETA* incorporates accessibility (ATAC-seq) and expression data (RNA-seq) to infer the gene targets of regulatory loci (ATAC-seq peaks) by calculating a rank product, akin to a p-value, for each gene (RPg). The RP for n genes is calculated as RPg =(Rgb/n)*(Rge/n) where adjusted p-values corresponding to RNA-seq differential expression for each gene (Rge) are ranked in ascending order and ATAC-seq peak scores (Rgb) ranked in descending order. RNA-seq differential expression results for all genes (*DESeq2*) and ATAC-seq peak scores (*MACS2*) were converted into *BETA*-compatible formats and passed as inputs for the program. A window of 500kb from the transcription start site (TSSs) was used to identify potential regulatory peaks and all differentially expressed genes for a given condition were considered in the analysis. The resulting output, consisting of RPs for all genes expressed in the RNA-seq dataset, was then passed into R for downstream interrogation. Significant RPs were demarcated as genes which exhibited an RP < 0.01 and *DESeq2* adj. p-value < 0.05. Venn Diagrams of overlap amongst significant RP genes were generated and visualized using *Limma*.^54^ All enrichment analyses and dot plot visualizations were performed using the *clusterProfiler* package.^55^ Heatmaps of gene expression data were generated using the *ComplexHeatmap* package.^53^ The *Sushi* package was applied to plot ATAC-seq and RNA-seq traces along the hg38 genome.^62^

### Western Blotting

At time of collection, cells were washed 1x with PBS and lysed in cold 1x RIPA buffer supplemented with Pierce® Protease and Phosphatase Inhibitor Tablets (Thermo) by incubating for 5 minutes at room temperature followed by gentle cell scraping. Lysates were then incubated on a thermoshaker set to 4C at 1,500 RPM for 10 minutes. Cytoskeletal and ECM debris were removed from the lysate via centrifugation for 15 minutes at 17,000xg. Bulk protein concentration of the resulting lysates was evaluated using the Pierce 660nm Assay system (Thermo). Lysates were mixed with 4x Laemmli buffer (Alfa Aesar) containing DTT (Acros) and heat-denatured at 95C for 5 minutes. Proteins were separated by SDS-PAGE, transferred to nitrocellulose membrane, and blocked in PBST + 5% milk (Lab Scientific) for 1hr at room temperature. Across all studies, blotting was performed using primary antibodies for: HOXA5 (1:200, mouse, sc-365784, Santa Cruz Biotech), α-Tubulin (1:5,000, rabbit, 2144, Cell Signaling Technology), pSer9-GSK3β (1:1,000, rabbit, D85E12, Cell Signaling Technology), and GSK3β (1:1,000, mouse, 3D10, Cell Signaling Technology). After overnight incubation in primary antibody, membranes were incubated with HRP-conjugated anti-rabbit IgG (1:10,000, 7074, Cell Signaling Technology) or anti-mouse IgG (1:10,000, 7076, Cell Signaling Technology) as appropriate. Membranes were developed using the SuperSignal West Femto Maximum Sensitivity Substrate kit (Thermo) and signal acquired using ECL technique. In all cases, following detection of target antigen, membranes were stripped once using Restore PLUS Western Blot Stripping Buffer (Thermo) and re-probed for the loading control. Band intensity was quantified using ImageJ (National Institutes of Health) gel analyzer tools, and technical replicates were pooled for statistical analysis.

### Cytoskeletal Inhibitor studies

iHLF cultures were serum starved overnight in 1% FBS DMEM medium prior to seeding. Cells were seeded at a density of 8,000 cells/cm^2^ in 5kPa PDMS hydrogel (Excellness) or TCP 6-well plates coated with 10μg/mL full length human Fn. Cells were maintained in 1% FBS DMEM and allowed to adhere overnight. After allowing for adhesion, cells were stimulated with 1% FBS DMEM containing 10μM p160ROCK inhibitor Y-27632 (Enzo) for 6hr or 1μM Latrunculin A (Cayman Chem) for 3hr. The vehicle control medium used was 1% FBS DMEM supplemented with 0.2% DMSO. All treatments were collected at a singular endpoint.

### SFK inhibitor studies

iHLF cultures were serum starved overnight in 1% FBS DMEM medium prior to seeding. Cells were seeded at a density of 8,000 cells/cm^2^ on 5kPa PDMS hydrogel (Excellness) or TCP 6-well plates coated with 10μg/mL full length human Fn. Cells were maintained in 1% FBS DMEM and allowed to adhere overnight. After allowing for adhesion, cells were stimulated with 1% FBS DMEM containing 20μM SRC inhibitor KB SRC 4 (Tocris) or 10μM pan-Src Family Kinase (SFK) inhibitor PP2 (Tocris) for periods of 3hr or 6hr, respectively. The vehicle control medium used was 1% FBS DMEM supplemented with 0.2% DMSO. All treatments were collected at a singular end point.

### Fibronectin fragment cultures

Synthesis of Fn^4G^ and Fn^9*10^ peptide fragments was performed as previously described.^42^ In preparation for functionalization with fragments, TCP, 5kPa PDMS hydrogel (Excellness), or 100kPa PDMS hydrogel (Excellness) surfaces were first plasma-etched in a plasma cleaner for 45s. Following treatment, surfaces were coated with freshly made 1% APTES solution in H_2_O and incubated at 60C for 90 min. Surfaces were washed 3x with diH_2_O and allowed to air dry. Silylated surfaces were maleimide activated by coating with a 2mg/mL solution of Sulfo-SMCC (Thermo) for 1hr at room temperature. In parallel to maleimide activation, stocks of Fn fragments containing a C-terminal cysteine included for site-specific conjugation were reduced with Pierce™ Immobilized TCEP Disulfide Reducing Gel (Thermo) to ensure the presence of NH-reactive monomers. Briefly, a 3:2 (v/v) solution of reducing gel to fragment peptide solution was incubated on a skewer for 1hr at room temperature. After brief centrifugation, the supernatant containing the reduced Fn fragment peptides was collected and diluted to a working concentration of 30μg/mL. The maleimide-activated surfaces were then incubated with reduced Fn fragment at 4C overnight with gentle shaking. Control surfaces were incubated with 10μg/mL full length human Fn in parallel. The next day the coated surfaces were washed and any nonreacted maleimide groups were blocked with a 10μg/mL L-cysteine (Thermo) solution for 1hr at room temperature with gentle shaking. Surfaces were again washed and blocked with a 1% BSA solution for 1hr at room temperature with gentle shaking. After final washes the surfaces were ready for cell culture. In all cases, iHLF cultures were serum starved overnight in 1% FBS DMEM medium prior to seeding. Cells were seeded on Fn^4G^, Fn^9*10^, or full-length Fn control substrates at a density of 8,000 cells/cm^2^ in 1% FBS DMEM. Cells were collected for downstream analysis after 24hr in culture.

### Statistical Analysis

GraphPad Prism was used to perform all statistical analysis of molecular data. All bar graphs were presented as mean ± S.D. Two-tailed: 1-way ANOVA, 2-way ANOVA, Brown-Forsythe and Welch ANOVA, and Kruskal-Wallis tests were employed to determine significance. Tests were determined based on data distribution, variance, and statistical hypotheses. Significant differences were considered when *p* < 0.05 (*); *p* < 0.01 (**); *p* < .001 (***); *p* < .0001 (****).

## Acknowledgements

We thank the University of Virginia Flow Cytometry Core Facility for assisting with cell sorting and flow cytometry procedures. We also thank the University of Virginia Genome Analysis and Technology core for their assistance with NGS library quality control prior to sequencing.

## Author Contributions

A.E.M. designed, conducted and analyzed all experiments presented in figures 2-6; P.H. designed and conducted experiments presented in figure 1; R.B. assisted with experiments; R.T.H. assisted with experiments and analysis; A.E.M and T.H.B wrote the manuscript with edits from R.T.H, D.A, and M.C.; T.H.B. and M.C. financially supported the research.

## Materials & Correspondence

Correspondence to Thomas H. Barker

## References

1. Wilson, M.S. & Wynn, T.A. Pulmonary fibrosis: pathogenesis, etiology and regulation. Mucosal Immunol 2, 103–121 (2009).

2. Ihn, H. Pathogenesis of fibrosis: role of TGF-beta and CTGF. Curr Opin Rheumatol 14, 681–685 (2002).

3. Kumar, A., Khandelwal, N., Malya, R., Reid, M.B. & Boriek, A.M. Loss of dystrophin causes aberrant mechanotransduction in skeletal muscle fibers. FASEB J 18, 102–113 (2004).

4. Jaalouk, D.E. & Lammerding, J. Mechanotransduction gone awry. Nat Rev Mol Cell Biol 10, 63–73 (2009).

5. Di-Luoffo, M., Ben-Meriem, Z., Lefebvre, P., Delarue, M. & Guillermet-Guibert, J. PI3K functions as a hub in mechanotransduction. Trends Biochem Sci 46, 878–888 (2021).

6. Du, J. et al. Extracellular matrix stiffness dictates Wnt expression through integrin pathway. Sci Rep 6, 20395 (2016).

7. Martineau, L.C. & Gardiner, P.F. Insight into skeletal muscle mechanotransduction: MAPK activation is quantitatively related to tension. J Appl Physiol (1985) 91, 693–702 (2001).

8. Younesi, F.S., Miller, A.E., Barker, T.H., Rossi, F.M.V. & Hinz, B. Fibroblast and myofibroblast activation in normal tissue repair and fibrosis. Nat Rev Mol Cell Biol 25, 617–638 (2024).

9. Saalbach, A. & Anderegg, U. Thy-1: more than a marker for mesenchymal stromal cells. FASEB J 33, 6689–6696 (2019).

10. Dominici, M. et al. Minimal criteria for defining multipotent mesenchymal stromal cells. The International Society for Cellular Therapy position statement. Cytotherapy 8, 315–317 (2006).

11. Hu, P., Leyton, L., Hagood, J.S. & Barker, T.H. Thy-1-Integrin Interactions in. Front Cell Dev Biol 10, 928510 (2022).

12. True, L.D. et al. CD90/THY1 is overexpressed in prostate cancer-associated fibroblasts and could serve as a cancer biomarker. Mod Pathol 23, 1346–1356 (2010).

13. Mayeux-Portas, V., File, S.E., Stewart, C.L. & Morris, R.J. Mice lacking the cell adhesion molecule Thy-1 fail to use socially transmitted cues to direct their choice of food. Curr Biol 10, 68–75 (2000).

14. Hagood, J.S. et al. Loss of fibroblast Thy-1 expression correlates with lung fibrogenesis. Am J Pathol 167, 365–379 (2005).

15. Fiore, V.F. et al. Conformational coupling of integrin and Thy-1 regulates Fyn priming and fibroblast mechanotransduction. J Cell Biol 211, 173–190 (2015).

16. Liu, X. et al. Thy-1 interaction with Fas in lipid rafts regulates fibroblast apoptosis and lung injury resolution. Lab Invest 97, 256–267 (2017).

17. Ripamonti, U. Developmental pathways of periodontal tissue regeneration: Developmental diversities of tooth morphogenesis do also map capacity of periodontal tissue regeneration? J Periodontal Res 54, 10–26 (2019).

18. Bielefeld, K.A., Amini-Nik, S. & Alman, B.A. Cutaneous wound healing: recruiting developmental pathways for regeneration. Cell Mol Life Sci 70, 2059–2081 (2013).

19. Stabler, C.T. & Morrisey, E.E. Developmental pathways in lung regeneration. Cell Tissue Res 367, 677–685 (2017).

20. Hrycaj, S.M. et al. Hox5 Genes Regulate the Wnt2/2b-Bmp4-Signaling Axis during Lung Development. Cell Rep 12, 903–912 (2015).

21. Aubin, J., Lemieux, M., Tremblay, M., Bérard, J. & Jeannotte, L. Early postnatal lethality in Hoxa-5 mutant mice is attributable to respiratory tract defects. Dev Biol 192, 432–445 (1997).

22. Kinkead, R. et al. Respiratory adaptations to lung morphological defects in adult mice lacking Hoxa5 gene function. Pediatr Res 56, 553–562 (2004).

23. Hrycaj, S.M. et al. genes direct elastin network formation during alveologenesis by regulating myofibroblast adhesion. Proc Natl Acad Sci U S A 115, E10605–E10614 (2018).

24. Hrycaj, S.M., Marty-Santos, L., Rasky, A.J., Lukacs, N.W. & Wellik, D.M. Loss of Hox5 function results in myofibroblast mislocalization and distal lung matrix defects during postnatal development. Sci China Life Sci 61, 1030–1038 (2018).

25. Li, M.H. et al. The Lung Elastin Matrix Undergoes Rapid Degradation Upon Adult Loss of. Front Cell Dev Biol 9, 767454 (2021).

26. Wang, C.C. et al. HOXA5 inhibits metastasis via regulating cytoskeletal remodelling and associates with prolonged survival in non-small-cell lung carcinoma. PLoS One 10, e0124191 (2015).

27. Liang, Y., Zhou, R., Fu, X., Wang, C. & Wang, D. HOXA5 counteracts the function of pathological scar-derived fibroblasts by partially activating p53 signaling. Cell Death Dis 12, 40 (2021).

28. Fiore, V.F., et al. αvβ3 Integrin drives fibroblast contraction and strain stiffening of soft provisional matrix during progressive fibrosis. JCI Insight 3 (2018).

29. Roca-Cusachs, P., Gauthier, N.C., Del Rio, A. & Sheetz, M.P. Clustering of alpha(5)beta(1) integrins determines adhesion strength whereas alpha(v)beta(3) and talin enable mechanotransduction. Proc Natl Acad Sci U S A 106, 16245–16250 (2009).

30. Wan, H. et al. Thy-1 depletion and integrin β3 upregulation-mediated PI3K-Akt-mTOR pathway activation inhibits lung fibroblast autophagy in lipopolysaccharide-induced pulmonary fibrosis. Lab Invest 99, 1636–1649 (2019).

31. Sanders, Y.Y. et al. Thy-1 promoter hypermethylation: a novel epigenetic pathogenic mechanism in pulmonary fibrosis. Am J Respir Cell Mol Biol 39, 610–618 (2008).

32. Chrzanowska-Wodnicka, M. & Burridge, K. Rho-stimulated contractility drives the formation of stress fibers and focal adhesions. J Cell Biol 133, 1403–1415 (1996).

33. Yu, L., Hébert, M.C. & Zhang, Y.E. TGF-beta receptor-activated p38 MAP kinase mediates Smad-independent TGF-beta responses. EMBO J 21, 3749–3759 (2002).

34. Fearing, B.V. et al. Mechanosensitive transcriptional coactivators MRTF-A and YAP/TAZ regulate nucleus pulposus cell phenotype through cell shape. FASEB J 33, 14022–14035 (2019).

35. Dupont, S. et al. Role of YAP/TAZ in mechanotransduction. Nature 474, 179–183 (2011).

36. Sisson, T.H. et al. Inhibition of myocardin-related transcription factor/serum response factor signaling decreases lung fibrosis and promotes mesenchymal cell apoptosis. Am J Pathol 185, 969–986 (2015).

37. Ligresti, G., et al. CBX5/G9a/H3K9me-mediated gene repression is essential to fibroblast activation during lung fibrosis. JCI Insight 5 (2019).

38. Midgley, A.C. et al. Hyaluronidase-2 Regulates RhoA Signaling, Myofibroblast Contractility, and Other Key Profibrotic Myofibroblast Functions. Am J Pathol 190, 1236–1255 (2020).

39. Dixon, J.R. et al. Topological domains in mammalian genomes identified by analysis of chromatin interactions. Nature 485, 376–380 (2012).

40. Mansisidor, A.R. & Risca, V.I. Chromatin accessibility: methods, mechanisms, and biological insights. Nucleus 13, 236–276 (2022).

41. Wang, S. et al. Target analysis by integration of transcriptome and ChIP-seq data with BETA. Nature Protocols 8, 2502–2515 (2013).

42. Cao, L. et al. Detection of an Integrin-Binding Mechanoswitch within Fibronectin during Tissue Formation and Fibrosis. ACS Nano 11, 7110–7117 (2017).

43. Schmidt, S. & Debant, A. Function and regulation of the Rho guanine nucleotide exchange factor Trio. Small GTPases 5, e29769 (2014).

44. Rinn, J.L. et al. A dermal HOX transcriptional program regulates site-specific epidermal fate. Genes Dev 22, 303–307 (2008).

45. Illig, R., Fritsch, H. & Schwarzer, C. Spatio-temporal expression of HOX genes in human hindgut development. Dev Dyn 242, 53–66 (2013).

46. Hutlet, B. et al. Systematic expression analysis of Hox genes at adulthood reveals novel patterns in the central nervous system. Brain Struct Funct 221, 1223–1243 (2016).

47. Nicola, T. et al. Loss of Thy-1 inhibits alveolar development in the newborn mouse lung. Am J Physiol Lung Cell Mol Physiol 296, L738–750 (2009).

48. Corces, M.R. et al. An improved ATAC-seq protocol reduces background and enables interrogation of frozen tissues. Nat Methods 14, 959–962 (2017).

49. Andrews, S. FastQC: a quality control tool for high throughput sequence data. Available online at: http://www.bioinformatics.babraham.ac.uk/projects/fastqc (2010).

50. Martin, M. Cutadapt removes adapter sequences from high-throughput sequencing reads. 2011 17, 3 (2011).

51. Pertea, M., Kim, D., Pertea, G.M., Leek, J.T. & Salzberg, S.L. Transcript-level expression analysis of RNA-seq experiments with HISAT, StringTie and Ballgown. Nat Protoc 11, 1650–1667 (2016).

52. Love, M.I., Huber, W. & Anders, S. Moderated estimation of fold change and dispersion for RNA-seq data with DESeq2. Genome Biol 15, 550 (2014).

53. Gu, Z., Eils, R. & Schlesner, M. Complex heatmaps reveal patterns and correlations in multidimensional genomic data. Bioinformatics 32, 2847–2849 (2016).

54. Ritchie, M.E. et al. limma powers differential expression analyses for RNA-sequencing and microarray studies. Nucleic Acids Res 43, e47 (2015).

55. Yu, G., Wang, L.G., Han, Y. & He, Q.Y. clusterProfiler: an R package for comparing biological themes among gene clusters. OMICS 16, 284–287 (2012).

56. Wang, Z., et al. BART: a transcription factor prediction tool with query gene sets or epigenomic profiles. Bioinformatics 34, 2867–2869 (2018).

57. Langmead, B. & Salzberg, S.L. Fast gapped-read alignment with Bowtie 2. Nat Methods 9, 357–359 (2012).

58. Feng, J., Liu, T., Qin, B., Zhang, Y. & Liu, X.S. Identifying ChIP-seq enrichment using MACS. Nat Protoc 7, 1728–1740 (2012).

59. Liao, Y., Smyth, G.K. & Shi, W. featureCounts: an efficient general purpose program for assigning sequence reads to genomic features. Bioinformatics 30, 923–930 (2014).

60. Heinz, S. et al. Simple combinations of lineage-determining transcription factors prime cis-regulatory elements required for macrophage and B cell identities. Mol Cell 38, 576–589 (2010).

61. Lawrence, M. et al. Software for computing and annotating genomic ranges. PLoS Comput Biol 9, e1003118 (2013).

62. Phanstiel, D.H., Boyle, A.P., Araya, C.L. & Snyder, M.P. Sushi.R: flexible, quantitative and integrative genomic visualizations for publication-quality multi-panel figures. Bioinformatics 30, 2808–2810 (2014).

63. Bailey, T.L., Johnson, J., Grant, C.E. & Noble, W.S. The MEME Suite. Nucleic Acids Res 43, W39–49 (2015).

