## Supplemental Figures 1-11 for "Fibroblast mechanoperception instructs pulmonary developmental and pattern specification gene expression programs"

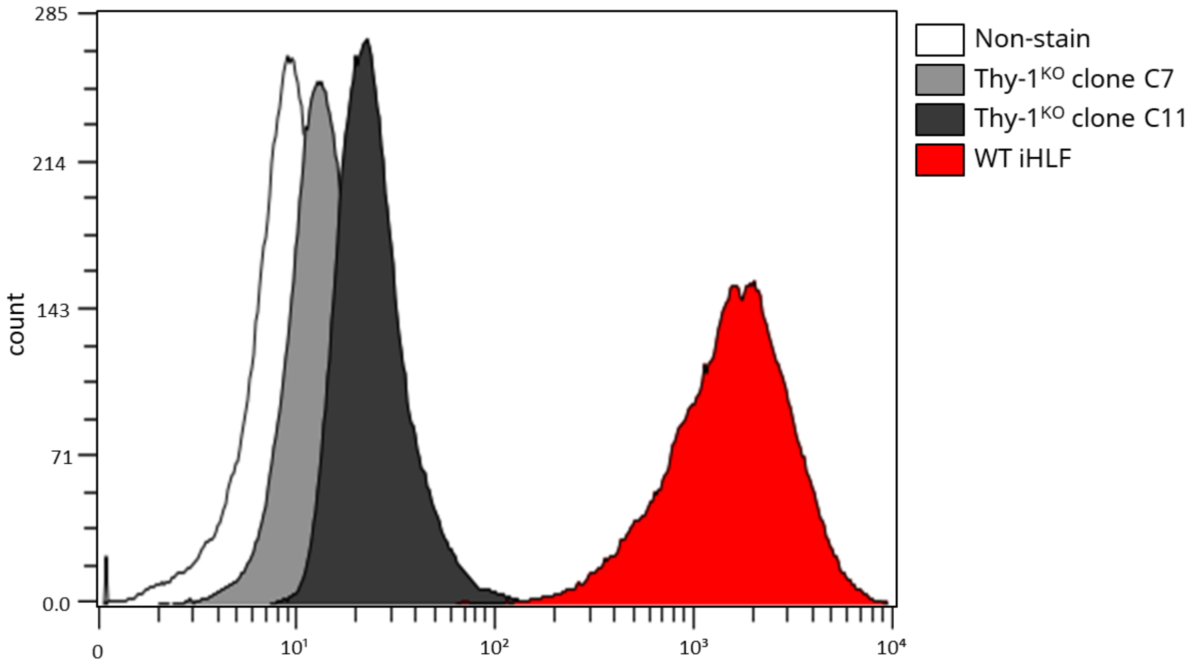


**Supplemental Figure 1:** Flow cytometry histograms for Thy-1 surface expression in immortalized human lung fibroblasts following CRISPR-Cas9 knockout of Thy-1 in clones C7 and C11 compared with WT and unstained control samples.

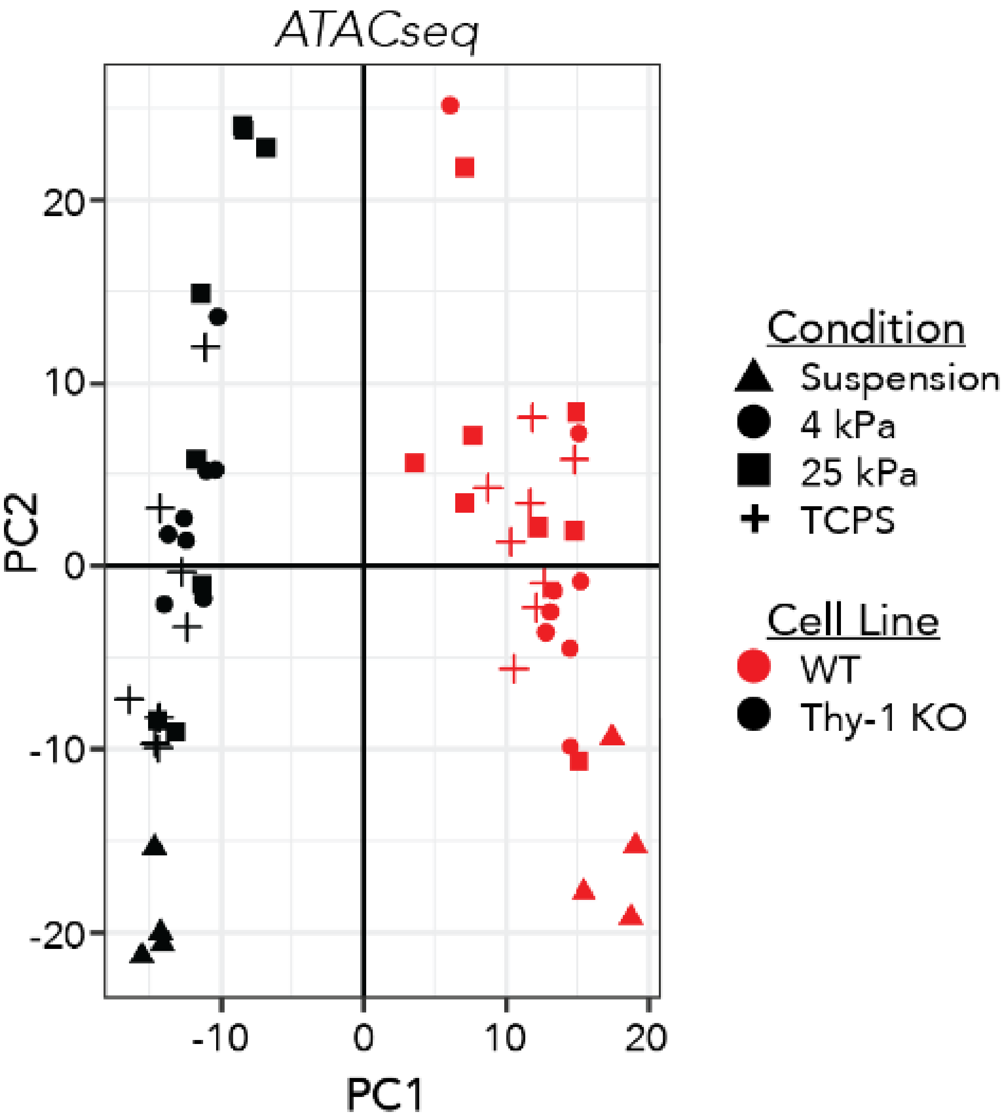


**Supplemental Figure 2:** Principal component analysis (PCA) of variance stabilized expression data based on weighted chromatin accessibility (loci) lists from ATAC-seq data. X and Y-axes represent the primary and secondary principal components (PC1, PC2), respectively. PC1= 48.20% of variance, PC2= 36.68% of variance.

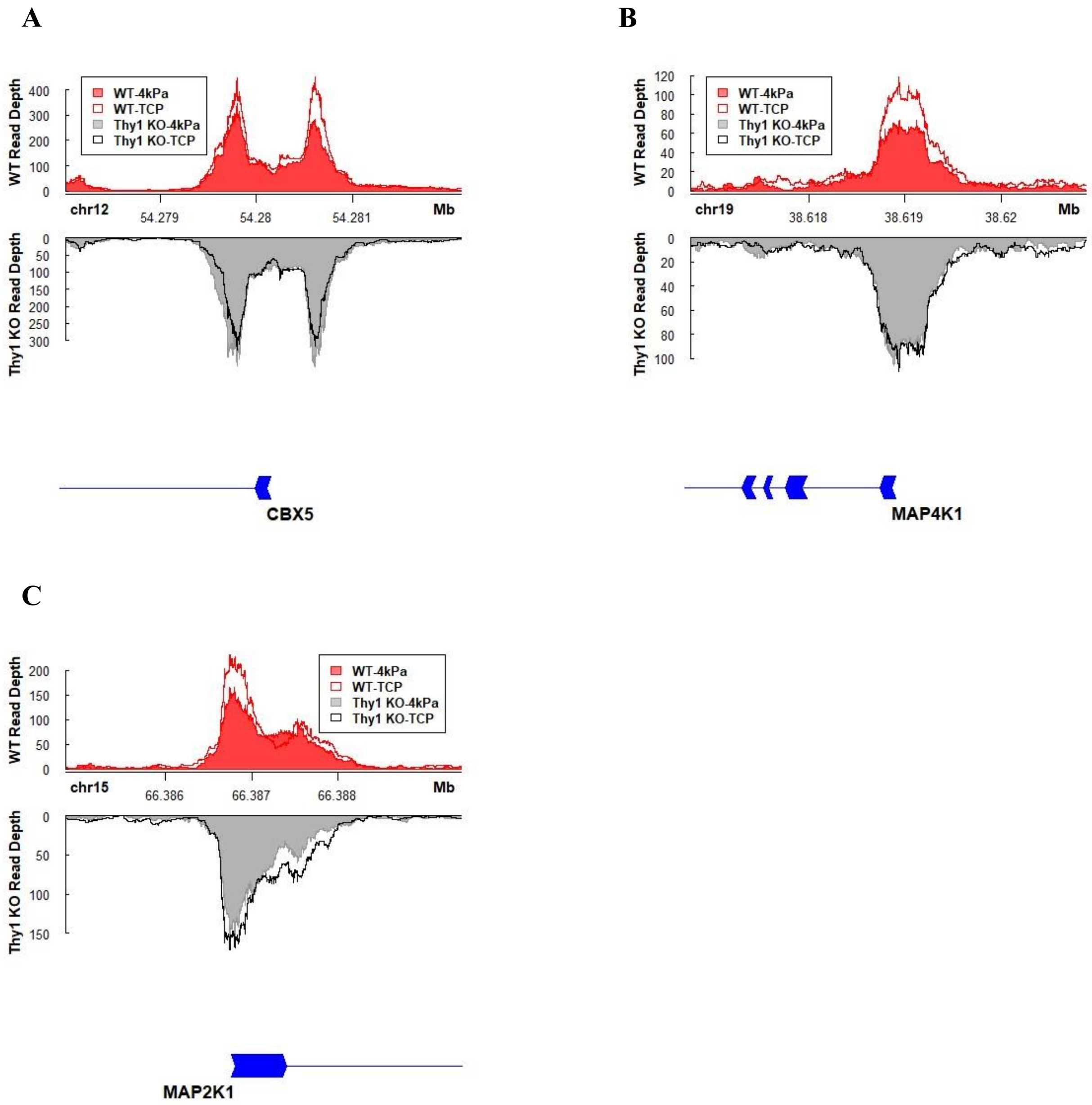


**Supplemental Figure 3:** Visualization of normalized ATAC-seq read pileup in WT (top; red) and Thy-$1^{KO}$ (bottom; black/grey) fibroblasts show deviations in promoter accessibility as a function of substrate stiffness at pro-fibrotic loci (A) *CBX5* (24hr), (B) *MAP4K1* (72hr), and (C) *MAP2K1*. Opaque traces represent 4kPa stiffness conditions; transparent traces represent 1GPa/TCP conditions.

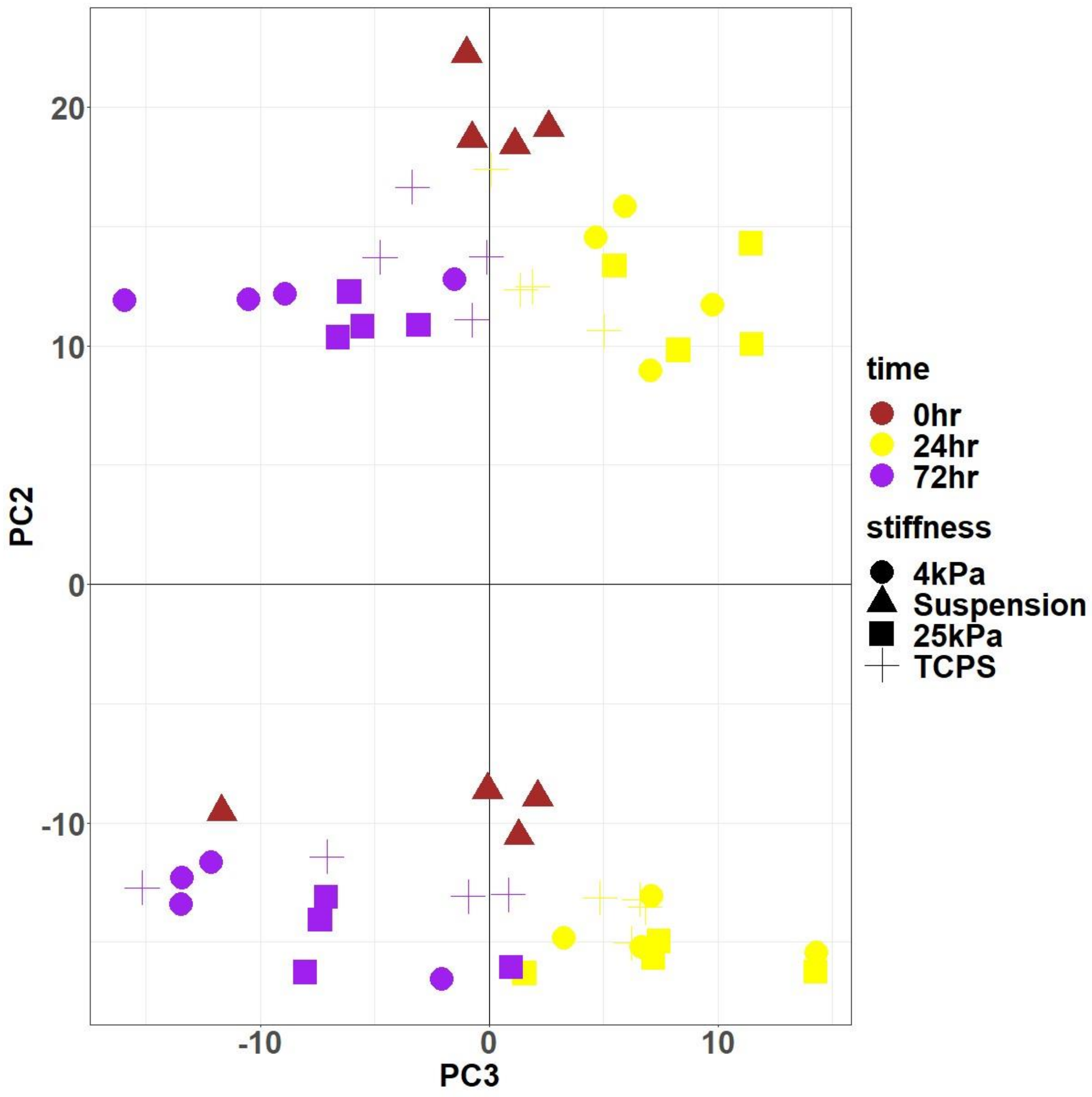


**Supplemental Figure 4:** Analysis of extended principal components from gene-loading data (RNA-seq) reveals PC3 (X-axis) separates gene expression data based on culture time. Top cluster represents WT samples while bottom cluster represents Thy-$1^{KO}$ samples. PC2= 36.68% of variance, PC3= 4.66% of variance.

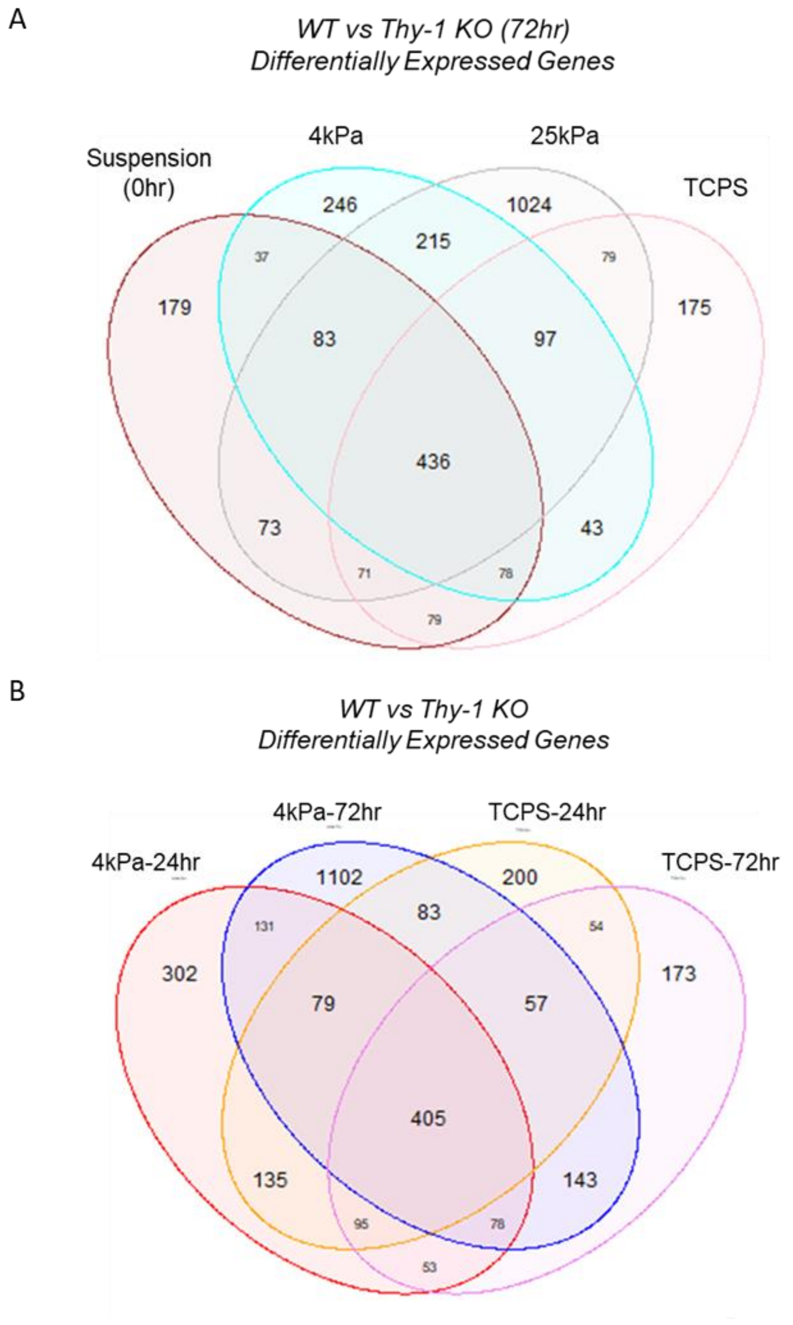


**Supplemental Figure 5:** Comparison of differentially expressed genes between WT and Thy-$1^{KO}$ populations (A) on all stiffnesses at 72hr in culture and (B) on 4kPa and TCPS substrates after 24hr or 72hr in culture. In all cases a conserved set of genes displayed significant changes in expression after Thy-1 loss (Wald test with BH correction; $p < 0.05$).

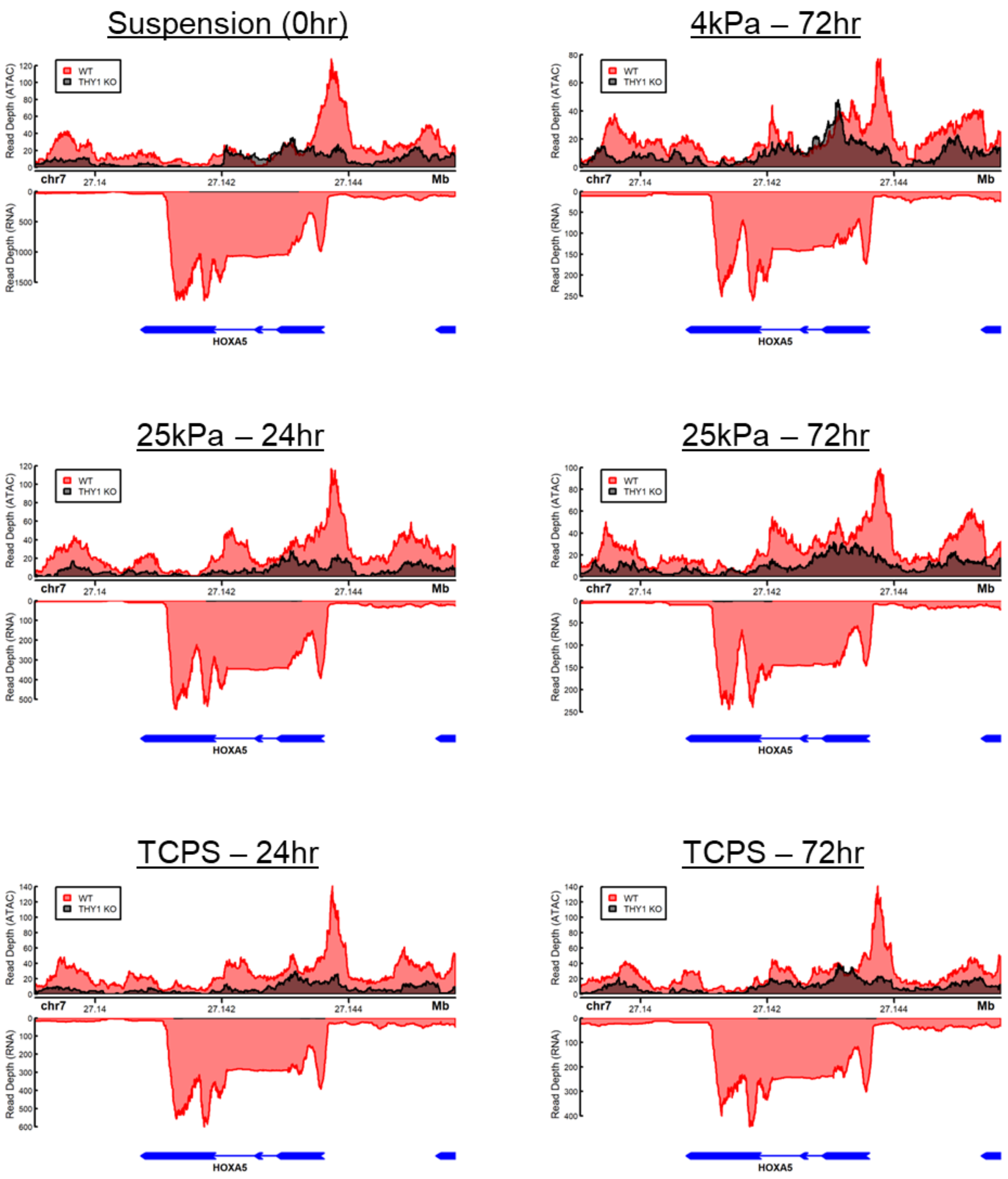


**Supplemental Figure 6:** Traces of read pileup in WT (red) and Thy-1$^{KO}$ (black) fibroblasts along the *HOXA5* gene body in each tested condition. Top panel represents ATAC-seq data; bottom panel represents RNA-seq data.

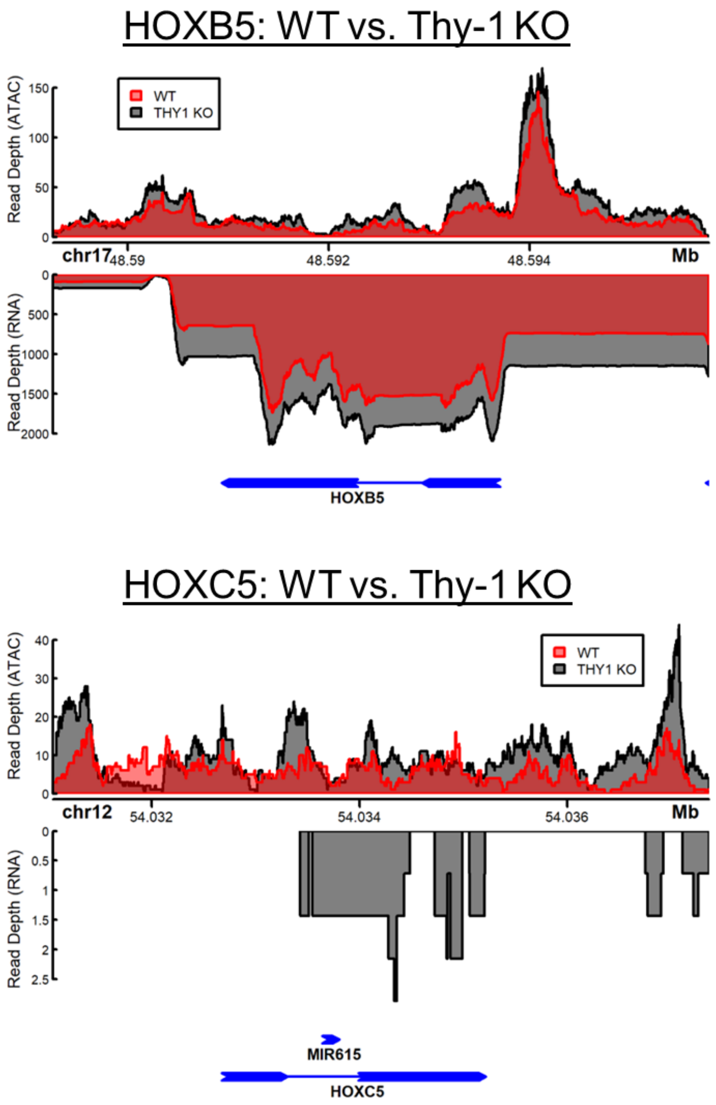


**Supplemental Figure 7:** Traces of read pileup in WT (red) and Thy-1$^{KO}$ (black) fibroblasts along the *HOXB5* and *HOXC5* gene bodies after culture on 4kPa substrates for 24-hours. Top panel represents ATAC-seq data; bottom panel represents RNA-seq data. No RNA-seq data is shown for WT sample at the *HOXC5* locus due to an absence of detectable RNA signal.

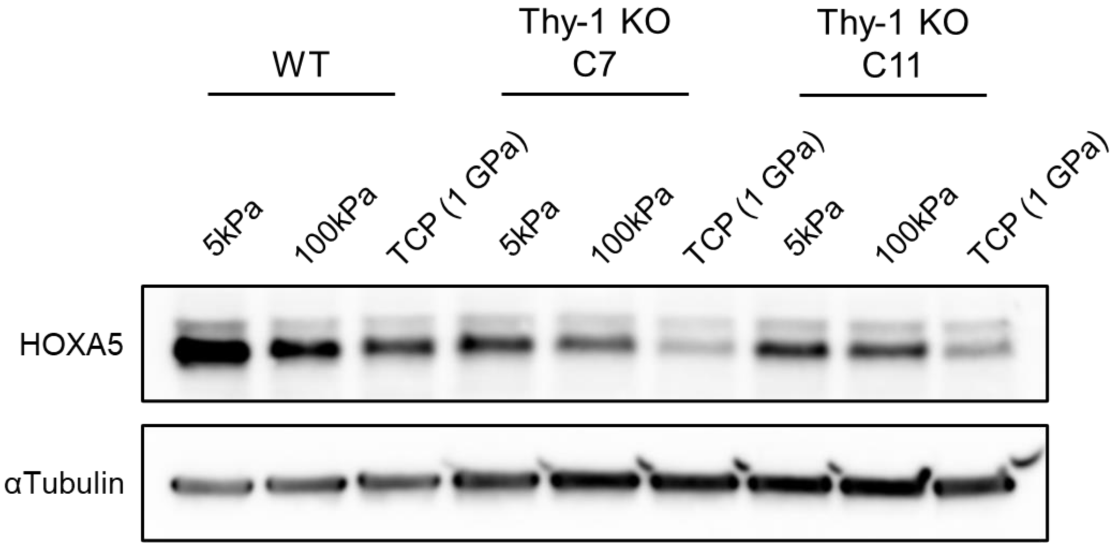


**Supplemental Figure 8:** Western blots for HOXA5 protein expression cultured on 5kPa, 100kPa, or tissue culture plastic (~1GPa) in WT fibroblasts and two independent Thy-$1^{KO}$ CRISPR-Cas9 clones.

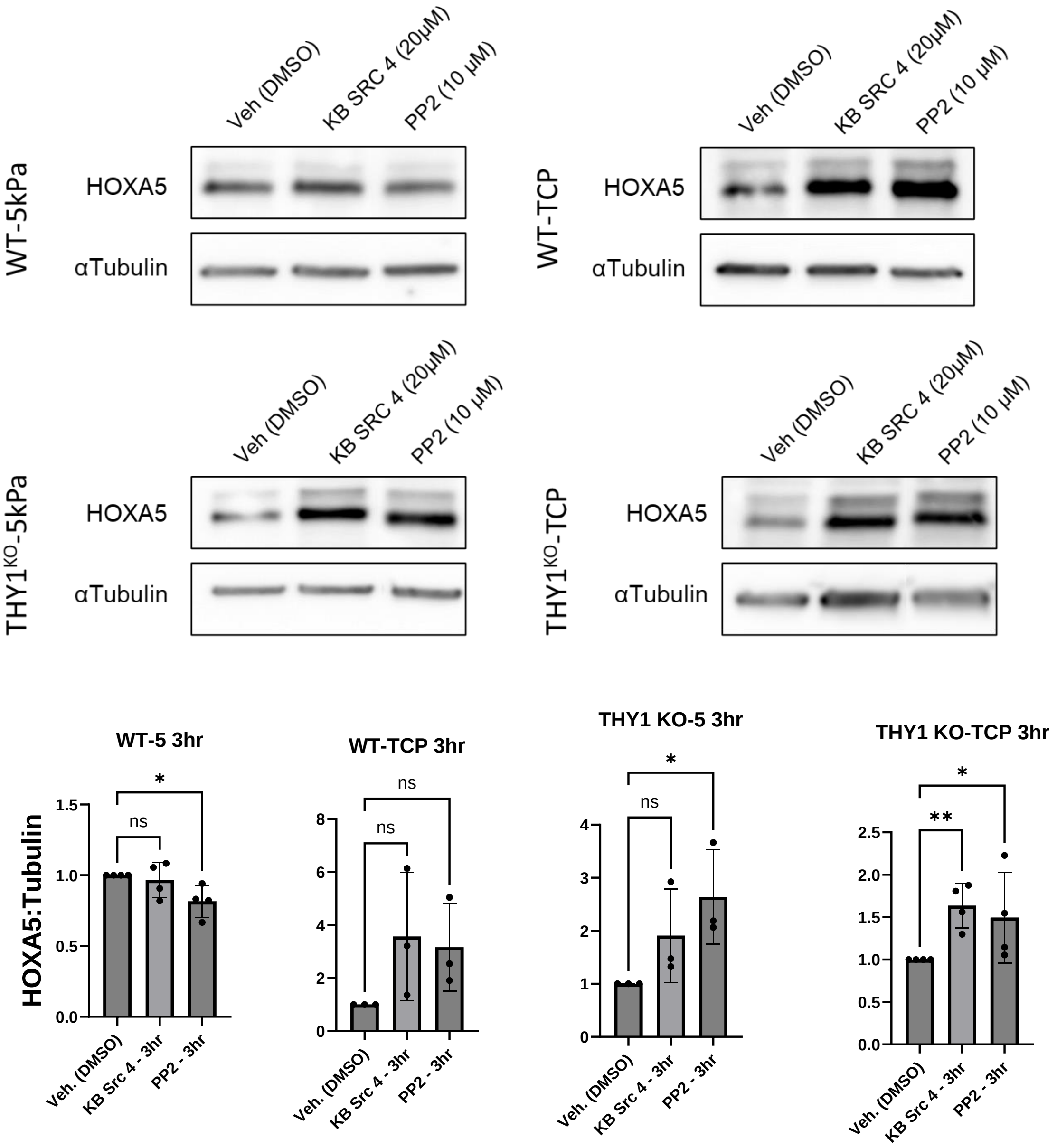


**Supplemental Figure 9:** Western blots for HOXA5 expression in WT and Thy-1$^{KO}$ fibroblasts cultured on soft hydrogels (4kPa) or stiff, tissue culture plastic (TCP; ~1GPa) substrates. Samples were allowed to adhere overnight then treated with 10µM pan-SRC family kinase (pan-SFK) inhibitor PP2 or 20µM SRC inhibitor KB SRC 4 for 3hr. Plots represent ratio of signal to vehicle condition after normalization of HOXA5 signal to loading control. Mean ± S.D. plotted; N=3 (Thy-1$^{KO}$-5kPa and WT-1GPa/TCP) or N=4 (WT-5kPa and Thy-1$^{KO}$-1GPa) independent experiments; WT-5kPa, Thy-1$^{KO}$ -5kPa and Thy-1$^{KO}$-1GPa = non-parametric Kruskal-Wallis test with post-hoc uncorrected Dunn's test and WT-1GPa = 1-way ANOVA with post-hoc uncorrected Fisher's LSD test, Tests chosen based on data distribution and variance. For all statistical tests: ns = $p > 0.05$; $p < 0.05$ (*); $p < 0.01$ (**); $p < .001$ (***); $p < .0001$ (****).

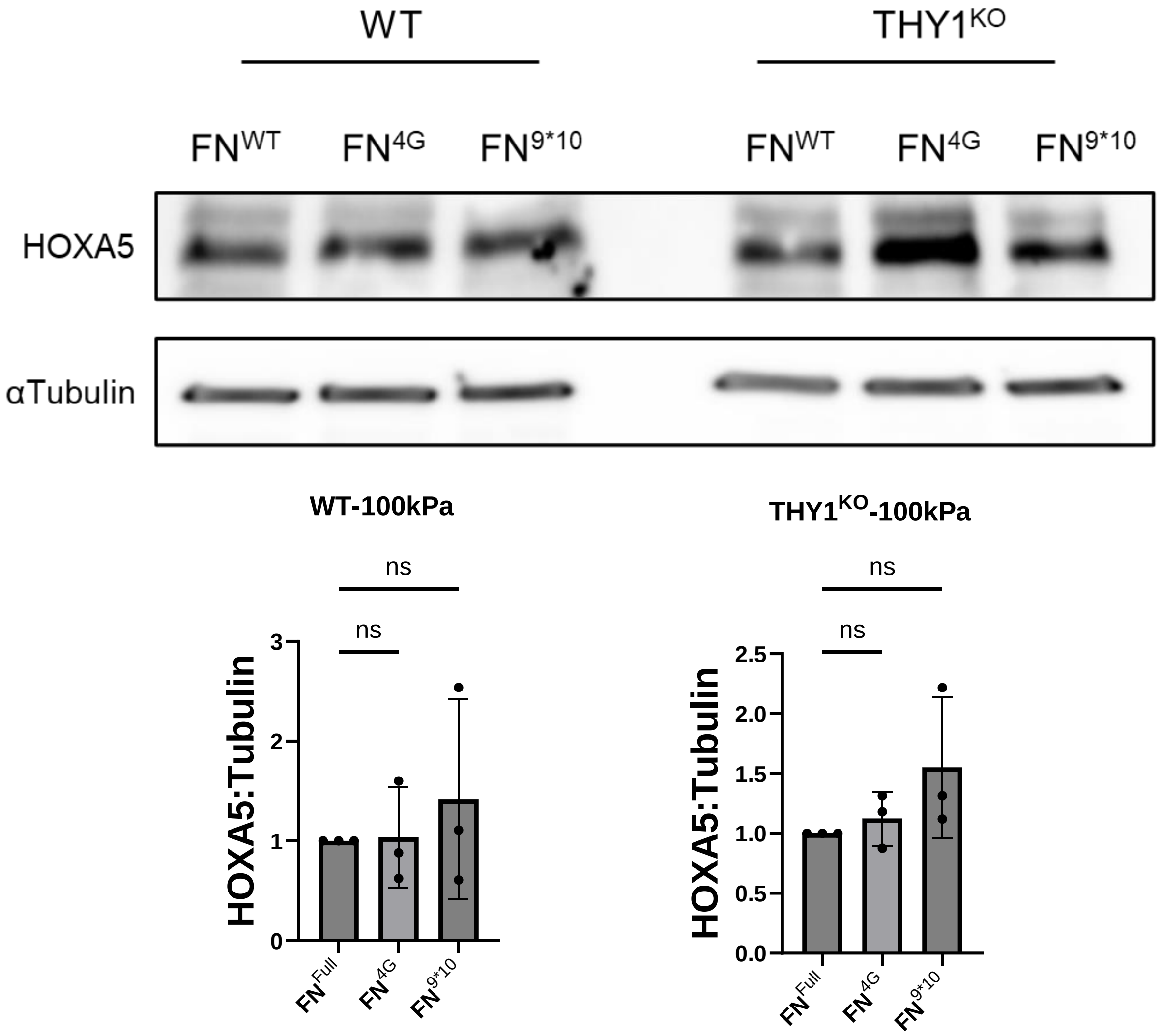


**Supplemental Figure 10:** Western blots for HOXA5 expression in WT and Thy-$1^{KO}$ fibroblasts cultured on 100kPa hydrogels coated with full-length (WT), 4G, or 9*10 fibronectin for 24 hours. No significant changes in HOXA5 expression were identified. Plots represent ratio of signal to full-length condition after normalization of HOXA5 signal to loading control. Mean ± S.D. plotted; N=3 independent experiments; WT-100kPa = 1-way ANOVA with post-hoc uncorrected Fisher's LSD test, Thy-$1^{KO}$-100kPa = Brown-Forsythe and Welch ANOVA with post-hoc unpaired t-tests with Welch's correction. For all statistical tests: ns = $p > 0.05$; $p < 0.05$ (*); $p < 0.01$ (**); $p < .001$ (***); $p < .0001$ (****).

### HOXA5

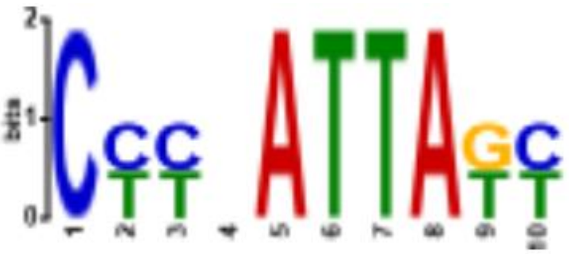

(Consensus Motif –CYYNATTAKY)

| Test Condition | Background Condition | *E*-value |
|---|---|---|
| WT-4kPa-24hr | Thy-1 KO-4kPa-24hr | **< 2.09e-11** |
| WT-TCP-24hr | Thy-1 KO-TCP-24hr | **< 1.59e-2** |

**Supplemental Figure 11:** Enrichment analysis for the HOXA5 consensus binding motif between WT and Thy-1 KO fibroblasts cultured on soft (4kPa) or stiff (TCP) substrates using *SEA* from the *MEME* family of packages. $E$ < 1e-3 was applied to determine a significant change in global binding site accessibility.

**Supplemental Tables (Provided as .xlsx Files)**

**Supplementary Table 1:** Gene-by-gene loadings for the top 50 transcripts that contributed the greatest degree of variance among the top 2 principal components from RNA-seq dataset.

**Supplementary Table 2 –** Binding Analysis for Regulation of Transcription (BART) outputs for displayed comparisons. The ten regulators with the lowest Irwin Hall p-values are presented for each comparison. For each comparison shown, differentially expressed genes (DEGs) downregulated in Thy-1 KO fibroblasts relative to WT samples were input into the BART package and analyzed for known, *cis* binding transcriptional regulators that were enriched above a scrambled control. Irwin Hall p-value < 1e-3 was used to denote significance.
